# Maladaptive oxidative stress cascade drives type I interferon hyperactivity in TNF activated macrophages promoting necrosis in murine tuberculosis granulomas

**DOI:** 10.1101/2020.12.14.422743

**Authors:** Eric Brownhill, Shivraj M. Yabaji, Vadim Zhernovkov, Oleksii S. Rukhlenko, Kerstin Seidel, Bidisha Bhattacharya, Sujoy Chatterjee, Hui A. Chen, Nicholas Crossland, William Bishai, Boris N. Kholodenko, Alexander Gimelbrant, Lester Kobzik, Igor Kramnik

**Author notes:** Contributed equally (Eric Brownhill, Shivraj M. Yabaji). Corresponding author Igor Kramnik, MD, PhD Boston University School of Medicine, NEIDL 620 Albany St., Boston, MA 02118.

## Abstract

Tuberculosis remains a critical infectious disease world-wide. The development of novel therapeutic strategies requires greater understanding of host factors that contribute to disease susceptibility. A major unknown in TB pathogenesis is the mechanism of necrosis in TB granulomas that leads to the massive lung tissue damage and cavity formation necessary for the pathogen transmission. In humans, TB progression has been linked to hyperactivity of type I IFN (IFN-I) pathway, the primary cause of which remains elusive.

We studied the mechanistic drivers of pulmonary TB progression using a unique model B6J.C3-*Sst1*^*C3HeB/Fej*^ Krmn mice that develop human-like necrotic TB granulomas and IFN-I hyperactivity. We established that IFNβ super-induction occurred in the susceptible macrophages in response to continuous TNF stimulation in the context of a dysregulated antioxidant defense. We observed that unresolving oxidative stress amplified the induction of IFNβ through JNK activation and induced the Integrated Stress Response via PKR activation as a compensatory pathway. Subsequently, PKR amplifies IFNβ upregulation, forming a positive feedback loop, maintaining the hyperinflammatory state in susceptible macrophages and leading to mitochondrial dysfunction. Thus, within the inflammatory milieu, a cell-intrinsic mechanism of chronic regulatory dysfunction and unresolved stress gradually weakens the macrophage and ultimately promotes the necrotization of TB granulomas. The aberrant macrophage response to TNF can be prevented by an iron chelator and inhibitor of lipid peroxidation, ferrostatin-1. Moreover, ferrostatin treatment increased macrophage survival and boosted bacterial control in the TNF-stimulated macrophages infected with virulent Mtb. These findings identify targets for host-directed therapeutics to interrupt necrotization in TB granulomas.

## INTRODUCTION

Thousands of years of co-evolution with modern humans has made *Mycobacterium tuberculosis* (Mtb) arguably the most successful human pathogen(1). It currently colonizes approximately a quarter of the global population(2). Mtb is a specialized pathogen - compared to environmental bacteria, it has lost a significant portion of its genome along with its environmental niche, becoming fully dependent on humans for its maintenance and spread(3). The evolutionary success of Mtb relies on the ability to infect humans via the respiratory route, establish chronic persistence and, in a minority of the infected individuals, destroy lung tissue to form cavitary lung lesions and ensure efficient transmission via infectious aerosol particles(4). Most people with persistent infection develop latent TB(5), in which the immune response sequesters the bacteria inside a granuloma structure. However, these granulomas typically develop necrotic centers comprised of cell debris, where the mycobacteria can survive and may grow. About 5-10% of latently infected individuals will eventually experience a failure of the granuloma structure and will develop active (and contagious) pulmonary TB(5, 6).

A major unanswered question in TB pathogenesis, which is central to understanding its evolutionary success, is the mechanism of necrosis in TB granulomas and the development of massive lung tissue damage with the formation of cavities (7). An important clue is that formation of large necrotic granulomas and cavities is observed only in a fraction of Mtb-infected individuals. Therefore, host factors play a major role in determining trajectories of the granulomas. Existing concepts explaining the host control of granuloma necrotization fall into two main categories: i) inadequate local immunity that allows exuberant bacterial replication and production of virulence factors(8) that drives tissue necrosis, and ii) excessive effector immunity that results in immune-mediated tissue damage(9). Although both scenarios are likely, they are mechanistically distinct and require different therapeutic strategies. Therefore, in-depth understanding of host-mediated mechanisms of necrosis in TB lesions is necessary for accurate patient stratification. It will also accelerate the development of effective means of immune modulation both at individual and population levels.

In a majority of humans, essential immune mechanisms at the whole organism level are intact and TB progression in about 85% of human TB is localized to the respiratory tract. Moreover, trajectories of individual lesions within a single host are often dissimilar, suggesting the importance of lesion-level control. This has sparked increased attention towards analyses of immune cell interactions within TB granulomas. A number of factors limit host immunity locally and transform granulomas into a protected niche harboring the pathogen and preventing its eradication. These include spatial separation of T cells and macrophages, inadequate immune cell turnover, formation of foamy macrophages, local excess of activating or immunosuppressive cytokines, hypoxia, angiogenesis and nutrient deprivation.

Mouse models that recapitulate various trajectories of the necrotic granuloma formation are necessary for in-depth mechanistic analyses of both the pathogen and host factors driving these processes, as well as for pre-clinical validation of precision “necrosis-directed” therapies. Although inbred mouse strains routinely used in TB research, such as C57BL/6 (B6) and BALB/c, do not develop necrotic TB granulomas, the necrotization of TB lesions can be recapitulated in experimental mouse models (reviewed in (10),(11)). Using forward genetic analysis, we have mapped several genetic loci of TB susceptibility. Surprisingly, a single locus on chromosome 1, *sst1* (***<u>s</u>***uper***<u>s</u>***usceptibility to ***<u>t</u>***uberculosis 1), was responsible for the control of the necrotization of TB lesions(12). This locus contains strong candidate genes Sp110b(13) and Sp140(14), whose expression is greatly diminished in mice that carry the *sst1* susceptibility allele. To study mechanisms of necrosis controlled by the *sst1* locus at cellular, tissue and whole organism levels, we made a congenic mouse strain B6J.C3-*Sst1C3HeB/Fej*Krmn (B6.Sst1S) that carries the C3HeB/FeJ-derived susceptibility allele of the *sst1* locus on the resistant B6 genetic background(15).

The B6.Sst1S mouse model recapitulates features of an ideal human host, from the pathogen’s evolutionary standpoint. (I) The animals are immunocompetent and do not rapidly succumb to disseminated infection. (II) Necrotic granulomatous lesions are formed exclusively in the lungs. (III) The necrotic granulomas are stable and separated from healthy lung tissue by well-organized multi-layer wall composed of fibrotic capsule and all major immune cell populations, preventing Mtb dissemination and severe lethal disease. Mechanistically, the *sst1* susceptible phenotype is associated with hyperactivity of the type I interferon (IFN-I) pathway in vitro and in vivo(16, 17). Hyperactivity of IFN-I pathway have also been found in humans with active TB (18), and an excess of IFNβ production in TB infection is known to be maladaptive on the part of the host macrophage (19).

By comparing responses of the inbred B6 wild type (B6wt) and B6.Sst1S mice to several intracellular bacterial pathogens in vivo and using reciprocal cell transfer experiments we found that the *sst1* locus primarily controls macrophage functions(13, 20, 21). Because of central importance of TNF in TB granulomas, we compared the kinetics of TNF responses of the WT and Sst1S macrophages in vitro. After prolonged stimulation with TNF, the Sst1S macrophages expressed higher levels of IFNβ and interferon-stimulated genes (ISGs) and upregulated markers of the integrated stress response (ISR) in an IFN-I -dependent manner. In parallel, they displayed evidence of protein aggregation and proteotoxic stress that could be inhibited by ROS scavengers and inhibitors of protein translation, but not by the type I IFN receptor blockade(16). Thus, the Sst1S macrophage activation displayed features of dysregulation of multiple stress and activation pathways controlled by a single genetic locus in a coordinated manner. These findings suggest that the *sst1*-encoded factors regulate a core response program that normally prevents pathological cascades that lead to TB progression.

To identify the specific molecular mechanisms that underlie the *sst1*-mediated necrotizing responses to Mtb, we have focused on cell-intrinsic factors that link an aberrant response to TNF and IFN-I pathway hyperactivity to cellular stress and macrophage damage. Herein, we described how B6.Sst1S macrophages respond inadequately to TNF stimulation by failing to induce key mechanisms to protect against reactive oxygen species (ROS). The resulting oxidative damage triggers cellular stress pathways, including JNK and PKR activation, which maintain the IFNβ pathway hyperactivity and the unresolving Integrated Stress Response that ultimately damage macrophages. We demonstrate that the dysregulated anti-oxidant response to TNF in macrophages prior their infection leads to persistent iron- and lipid peroxidation-mediated oxidative damage and render them susceptible to subsequent infection with intracellular bacteria. We propose that this maladaptive response leads to local damage in a context of TB granulomas and, eventually, to TB progression in immunocompetent hosts.

## RESULTS

### Enhanced susceptibility to necrotizing TB granulomas in Sst1S mutants linked to decreased macrophage resilience to chronic stimulation with TNF

After aerosol infection with 25 - 50 CFU of virulent Mtb H37Rv, the B6.Sst1S mice develop heterogeneous pulmonary lesions. Necrotic pulmonary lesions are typically observed during the third month of infection. As demonstrated in **Fig.1A**., multiple areas of focal granulomatous bronchopneumonia, resembling solid TB lesions in the parental *sst1*-resistant B6 mice (**Fig. 1C**), coexist with large, well-organized, caseating necrotic granulomas (**Fig.1D**). Importantly, no necrotizing granulomas are observed in other organs (15). In contrast to reports that described TB progression in the *sst1*-susceptible parental C3HeB/FeJ mice, the B6.Sst1S TB lesions are more heterogeneous, and are not uniformly necrotic. At this stage, we observed necrotizing pneumonia in less than 10% of the animals (**Fig.1B and E**). The development of necrotic pulmonary granulomas in the presence of lesions typical for immunocompetent, resistant hosts (i.e. C57BL/6; B6.WT), in combination with the chronic course of the disease, clearly demonstrate that the *sst1S*-mediated necrotization of pulmonary TB lesions is not due to systemic failure of an essential host resistance mechanism. Because the *sst1*-mediated intra-granulomatous necrosis occurs in a lesion-specific manner, it is more likely driven by local factors, leading to a gradual deterioration of the granuloma’s cellular constituents.

**Figure 1.**
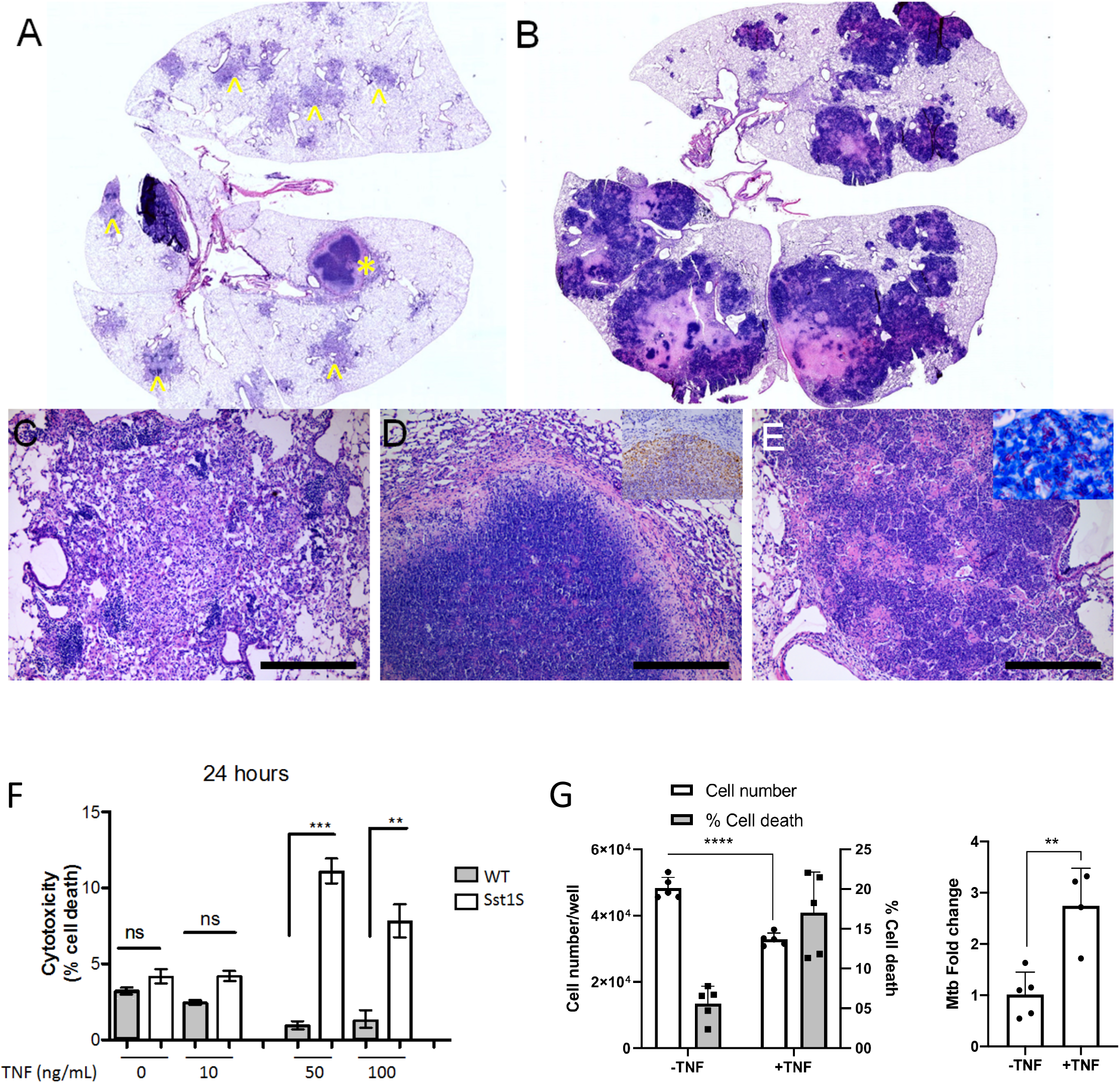
The Response to TB infection is a chronic and varied process involving the formation of progressive necrotic lesions. Histomorphologic phenotypes affiliated with chronic Mtb infection in B6.Sst1S mouse strain. A. Subgross image displaying heterogeneity of lesion size and type, including variably sized granulomatous bronchopneumonia (^) and a discrete caseating granuloma (*). Higher magnifications of the aforementioned lesions are depicted in Figures C and D respectively. B. Subgross image of necrotizing pneumonia phenotype. Although similar in histomorphologic features, there is diverse variability in lesion size. Higher magnification of this phenotype is outlined in Figure E. C. Focal granulomatous bronchopneumonia. Alveolar spaces adjacent to multiple bronchioles are flooded with macrophages, admixed with multifocal lymphoid aggregates. D. Caseating granuloma phenotype. Well demarcated, with a peripheral rim of collagen, radiating aggregate of macrophages, and central core of necrotic cellular debris. Inset. Immunohistochemistry (IHC). Abundant *Mycobacterium tuberculosis* antigen in the caseating granuloma, predominating within macrophages, but also within the necrotic core. E. Necrotizing pneumonia phenotype. Discrete focus of lytic necrosis consisting of necrotic cellular debris, fibrin, and degenerative neutrophils. Inset. Acid Fast Bacteria (AFB)-Ziehl-Neelsen Stain, abundant AFB within necrotic focus. Scale bars = 350 micrometers. A-C, 100x; Inset B, 630x; Inset C, 200x. F. Cell death in B6.Sst1S and B6.WT BMDMs stimulated with increasing concentrations of TNF for 24h. Cytotoxicity was measured by PI/Hoechst staining and automated microscopy. Representative experiment of 3 repeats. Error bars indicate standard deviation of technical replicates, significance values determined by paired t-test of technical replicates. G. B6.Sst1S macrophages were pretreated with TNFα (10 ng/mL) for 16 h or left untreated, then infected with Mtb at MOI 1. The cells were harvested 3 days post-infection and analyzed for cell survival and death. The Mtb load was observed using a qPCR-based method, normalized to a BCG spike for internal control. Error bars indicate standard deviation of technical replicates, significance values determined by 2-way ANOVA with multiple comparisons (cell death and survival) or unpaired t-test (Mtb fold change).

TNF plays major roles in the granuloma formation and maintenance both in human TB and in animal models (22). It is produced by myeloid cells and Mtb-specific T cells within TB granulomas. We have previously demonstrated that the B6.Sst1S macrophages develop an aberrant response to TNF in vitro within 12 - 18 h of TNF stimulation. This is characterized by upregulation of stress markers, but no cell death (16). To determine whether prolonged TNF stimulation (as likely present in the face of persistent mycobacterial infection) would be sufficient to induce death of the *sst1*-susceptible macrophages, we stimulated bone marrow-derived macrophages (BMDMs) isolated from the B6.WT and B6.Sst1S mice with increasing concentrations of TNF for 24 – 48 h. We observed that 50 or 100 ng/mL TNF induces a moderate increase in cell death in B6.Sst1S, while B6.WT macrophages remain resilient (**Fig.1F**). Next we tested whether this modest increase in cell death can be prevented by standard apoptosis or necroptosis inhibitors - pan-caspase inhibitor z-VAD and Nec-1, respectively. Neither of the inhibitors reduced the percentage of dead cells in TNF-treated B6.Sst1S BMDMs, nor eliminated the difference in cell death between TNF-stimulated B6 and B6.Sst1S BMDMs (**Suppl.Fig.1A and 1B**). Thus, increased death in the B6.Sst1S did not amount to the rapid and massive cell death induced by TNF in the presence of protein translation inhibitor cycloheximide, as in a necroptosis model (23), nor was it mediated by caspase activation.

We postulated that the *sst1*-mediated phenotype reflects chronic un-resolving stress which only modestly increases the probability of cell death per se, but which decreases the Sst1S macrophage resilience to pathogen-induced stress and impairs mycobacterial control. To test this hypothesis, we monitored survival and death rates of B6.Sst1S macrophages infected with mycobacteria, as well as Mtb growth rates, after 3 days with and without TNF stimulation. We observed that TNF stimulation reduced macrophage cell counts, and increased their death rates. The Mtb loads were also increased in TNF-treated B6.Sst1S macrophages (**Fig.1G**). We also compared the macrophage death between Sst1S and WT macrophages during a chronic infection course at low multiplicity of infection (MOI < 0.1). Use of low MOI allowed monitoring of the cells for longer periods and is expected to better model chronic disease in vivo. Treatment with 10 ng/mL TNF resulted in decreased survival of the B6.Sst1S BMDMs by day 8 of infection (**Suppl.Fig.1C and 1D**). Thus, in this in vitro model of Mtb infection, the discrimination between the WT and Sst1S phenotypes required chronic infection with virulent Mtb and prolonged exposure to TNF. These observations are consistent with a postulate that the susceptible macrophage phenotype emerges gradually as a result of an aberrant response to a combination of chronic inflammatory stimuli and bacterial infection.

### Unresolving stress underlies the aberrant response of the Sst1S macrophages to TNF

#### Dominant role of persistent TNF stimulation in the escalating IFN-I response

To determine the mechanisms of chronic, TNF-induced cell dysfunction in Sst1S macrophages, we initially focused on the IFN-I pathway, which is upregulated in Sst1S BMDMs and instrumental in escalation of the integrated stress response at early time points after TNF stimulation (16). We sought to determine whether the Sst1S macrophages were more sensitive than WT to chronic TNF stimulation in terms of the type I IFN pathway upregulation. To exclude effects of other soluble mediators produced by TNF-stimulated macrophages, we pre-stimulated BMDMs with TNF for 24 h, washed, replaced the medium, and either re-stimulated them with the same dose of TNF or left them untreated (**Fig.2A**). This experimental design allowed us to compare effects of the *sst1* locus on primary and secondary macrophage responses to TNF (**Fig.2A**, samples 1,2 and 3,4, respectively), as well as residual levels of transcripts that were maintained after the first 24 h TNF treatment in the absence of the second TNF stimulation (**Fig. 2A**, samples 5,6).

**Figure 2.**
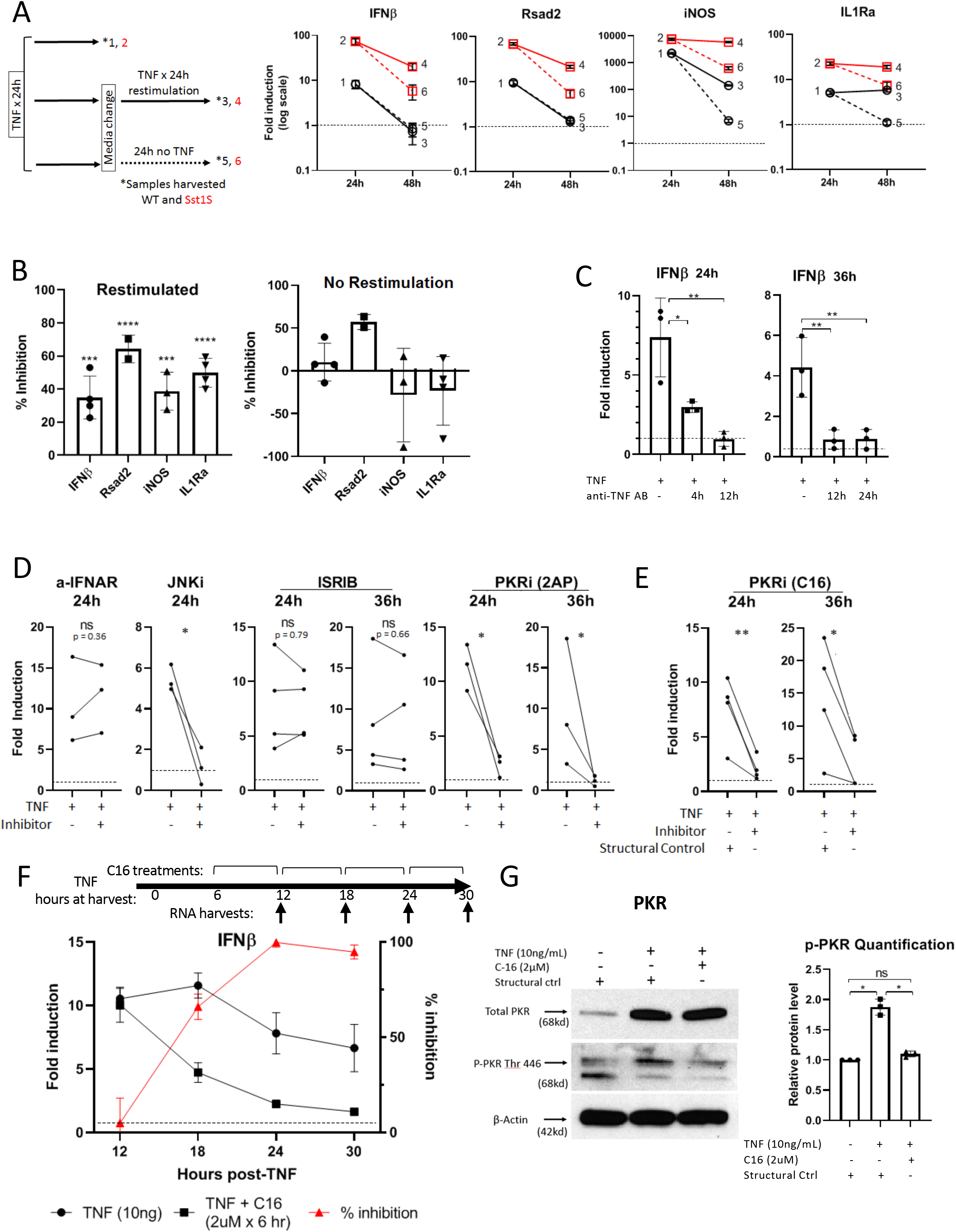
TNF induces and sustains a severe and prolonged IFNβ response in combination with JNK and PKR. A. Sst1S and WT BMDMs were stimulated with 10 ng/mL TNF for 24h, after which media was changed and cells were either re-stimulated with TNF or left unstimulated. Samples were harvested and qRT-PCR was performed after the first 24 h of stimulation, or 48 h after the original stimulation. One representative experiment shown out of four performed, error bars represent SD of technical replicates. Significance values in supplemental table 1 (two-way ANOVA). B. Sst1S BMDMs were treated as in (A), except upon media change at 24 h, were also treated with anti-IFNAR antibody or isotype control. Fold induction between antibody and control were compared to determine percent inhibition. Significance determined by comparison to isotype control (one-way ANOVA). C. Sst1S BMDMs were stimulated with 10 ng/mL TNF for 24 or 36 h or left unstimulated, and were treated with a TNF-blocking antibody 4, 12, or 24 h before harvest, as indicated, or left untreated as a control. qRT-PCR was performed on sample RNA to determine IFNβ induction, compared to untreated samples (dotted line). Significance determined by RM-ANOVA with multiple comparisons. D. Sst1S BMDMs were treated with TNF and analyzed by qRT-PCR as in (C), but inhibitors were added 6 h before harvest to determine the dependence of IFN on pathways downstream of TNF stimulation. Inhibitors include anti-IFNAR antibody, JNK inhibitor sp600125, Integrated Stress Response inhibitor (ISRIB), and the PKR inhibitor 2-aminopurine. E. Dependence of IFN on PKR was confirmed by use of C16, a more specific PKR inhibitor, in conjunction with its structural control. Significance determined by ratio-paired t-tests. F. Representative time-course of C16’s effect on IFN induction from 12 to 30 h of TNF treatment in Sst1S. C16 added 6 h before each harvest point. Fold induction of TNF + c16 compared to TNF only and untreated to determine % inhibition by c16. Error bars represent SD of technical replicates. G. Left: Example Western blot of Thr 446 Phospho- and total PKR with actin loading control in Sst1S. PKR is both induced and activated in Sst1S BMDMs by TNF alone at 24 h. Right: Quantification of three western blots confirming PKR activation. Densitometry by ImageJ software. Significance by RM-ANOVA with multiple comparisons. Statistical analysis: *p < 0.05, ** p < 0.01, *** p < 0.001, **** p <0.0001

In agreement with our previous studies, after the first 24 h TNF stimulation, we observed significantly higher upregulation of IFNβ and several inflammatory genes, including known targets of IFN-I in Sst1S BMDMs, as compared to the WT (**Fig.2A**, samples 1 and 2). The expression of IFNβ and Rsad2 mRNAs in the WT macrophages were refractory to the second TNF stimulation, as both IFNβ and Rsad2 levels returned to baseline at 48h either in the presence of in the absence of TNF (**Fig.2A**, samples 3 and 5, respectively). In contrast, the Sst1S macrophages (i) responded to the second TNF stimulation and (ii) expressed elevated IFNβ mRNA levels even in the absence of TNF re-stimulation (samples 4 and 6, respectively). Interestingly the level of IFNβ expression in the non-restimulated Sst1S macrophages was maintained at a similar level to that seen in WT after primary stimulation with TNF for 24 h (samples 6 and 3, respectively). Similar expression patterns were observed for a well-known IFN-I responsive gene, Rsad2. Distinct patterns of expression were observed for iNOS and IL1Ra, whose upregulation is also mediated by *sst1*. These genes responded to the second round of TNF stimulation in both B6.WT and B6.Sst1S macrophages (samples 3 vs 5 and 4 vs 6, respectively). However, their expression levels after TNF re-stimulation (at 48 h) in the B6.WT were similar to those of the Sst1S macrophages without re-stimulation (samples 3 and 6, respectively). These findings suggest that the Sst1S macrophages either lack negative feedback mechanisms of transcriptional regulation for IFNβ and IFN-regulated genes, or activate additional positive feedback and feed-forward circuits that maintain the IFN-I pathway upregulation.

To establish the extent to which this Sst1S-specific phenotype was dependent on IFNβ signaling through the interferon-alpha receptor (IFNAR), a known feed-forward element in the IFN-I signaling pathway, we added anti-IFNAR antibody along with the media change at 24 hours (h), to block further signaling through IFNAR after this point. The IFNAR1 blockade partially reduced the secondary TNF responses (**Fig.2B, left panel**). However, its effect was modest, especially in the case of IFNβ and iNOS transcripts. In addition, the residual expression of IFNβ, iNOS and IL1Ra (which are observed in TNF pre-stimulated Sst1S BMDMs in the absence of TNF re-stimulation) was unaffected by the IFNAR blockade (**Fig.2B, right pane**l). In contrast, administering a TNF-blocking antibody at 12 or at 24 h after stimulation with TNF resulted in reduction of IFNβ transcript levels to near-baseline both at 24 and 36 h (**Fig. 2C**). Hence, persistent TNF, but not IFN-I, signaling is necessary for sustaining the IFN-I pathway upregulation in B6.Sst1S macrophages at 24 – 36 h, as well as during the earlier 12 – 24 h period of TNF stimulation (**Fig.2C, left panel**). These data implicated the *sst1* locus in feedback regulation of TNF responses upstream of the IFN-I pathway escalation.

#### JNK and PKR activation downstream from TNF sustain IFN-I hyperactivity

Previously, we determined that the initial superinduction of IFNβ from 12 – 16 h of TNF stimulation was due to synergistic effects of NF-κB- and JNK-mediated pathways (16). Next, we wanted to explore pathways involved in sustaining the elevated IFNβ expression in Sst1S macrophages downstream of TNF signaling. We extended our previous findings by demonstrating that the JNK inhibitor SP600125, added 18 h after TNF was effective at inhibiting IFNβ levels by 24 h (**Fig.2D**). This inhibitor was also efficient when added during the 30 – 34 h interval (**Supplemental Fig.2B**). Thus, this stress-activated kinase plays a central role both in promoting the IFNβ superinduction within 8 – 16 h of TNF stimulation, as well as in sustaining the IFNβ upregulation at later time points in TNF-stimulated Sst1S macrophages.

Our observations of elevated IFNβ expression in TNF-primed Sst1S macrophages even after TNF withdrawal (**Fig.2A**) prompted us to search for additional mechanisms involved in sustaining the IFNβ upregulation in the susceptible macrophages. We hypothesized that an unresolving integrated stress response (ISR) may drive JNK activation at later stages, because prolonged translation inhibition is known to induce ribotoxic stress and JNK activation (24). However, a universal ISR inhibitor ISRIB that efficiently blocks ISR in our model (16), had no effect on the IFNβ expression during the 24 – 36 period. Surprisingly, the PKR inhibitor 2-aminopurine did suppress the IFNβ induction by TNF during this interval (**Fig.2D**). We confirmed this unexpected result using another PKR inhibitor, C16. The PKR specificity of the C16 effect on the IFNβ upregulation was confirmed using an inactive structural control (**Fig. 2E**). This observation was especially puzzling because no role for PKR in IFNβ induction was shown at an earlier step of the IFNβ superinduction, between 12 and 16 h in our previous studies (16).

To explore this apparent contradiction, we studied the effects of PKR inhibition with C16 on IFNβ expression at various stages of TNF stimulation more precisely: the inhibitor was added at 6, 12, 18 and 24 h after TNF stimulation and RNA samples were collected 6 h after each treatment (**Fig.2F diagram**). We confirmed that PKR inhibition had no effect on IFNβ upregulation at the initial 6 - 12 h interval but observed partial inhibition at 12 - 18 h and near-complete suppression at 24 and 30 h (**Fig.2F**). We confirmed PKR upregulation and activation in TNF-stimulated B6.Sst1S BMDMs during this period using western blot with total and activated (Thr 446 Phospho-PKR) PKR-specific antibodies (**Fig.2G**). Thus, the PKR contribution to IFNβ upregulation gradually increased after 18 h of TNF stimulation. However, this effect could not be explained by a PKR-mediated ISR, because it was not blocked by ISRIB. Nevertheless, alternative targets of activated PKR are known, including apoptosis stimulating kinase ASK1 (MAP3K5), which is upstream of JNK1 (25). Thus, the increased role of PKR in IFNβ maintenance, perhaps by engaging additional targets, signifies transition from the early “super-induction” (12 – 18 h) to a later “maintenance” (24 −48h) stage of the B6.Sst1S TNF response. Theoretically this transition permits the formation of a positive feedback loop, such as JNK ➢ IFNβ ➢PKR ➢JNK, maintaining and potentially amplifying the aberrant TNF response.

#### PKR limits the ISR escalation caused by prolonged TNF stimulation

In our previous experiments, we observed rapid upregulation of the ISR markers Trb3 and Chac1 in Sst1S BMDMs between 12 and 16 h of TNF stimulation. Their abrupt induction was mediated by IFN-I, PKR and ISR signaling (16). To monitor the ISR status during the “maintenance” phase, we examined Trb3 and Chac1 mRNA expression during 24 – 48 h of TNF stimulation. Trib3 and Chac1 mRNAs were highly expressed in TNF stimulated Sst1S BMDMs at 24 h, but not at all in WT (**Fig.3A**, samples 1 and 2). Next, we compared the mRNA levels in cells that were either re-stimulated with TNF for an additional 24 h after the initial TNF treatment, or not re-stimulated (as depicted in diagram in **Fig.2A**). The WT cells showed no upregulation of the ISR genes after TNF re-stimulation (samples 3 and 5), as well. In contrast, the ISR upregulation was sustained in the Sst1S macrophages to 48 h with or without TNF re-stimulation (**Fig. 3A**, samples 4 and 6). Their levels, however, were higher in the presence of TNF. This upregulation was partially dependent on IFNAR signaling (**Fig.3B**, left panel). Interestingly, the elevated expression of the ISR markers in the absence of TNF re-stimulation were maintained in the presence of IFNAR blocking antibodies (**Fig.3B**, right panel). In contrast, TNF blockade 12 h before harvest significantly reduced Trib3 levels at 24 and 36 h (**Fig. 3C**), i.e. the ISR upregulation required persistent TNF signaling, resembling an expression pattern of IFNβ (**Fig.2B**). We hypothesized that PKR activity may be involved in the ISR maintenance, as well.

**Figure 3.**
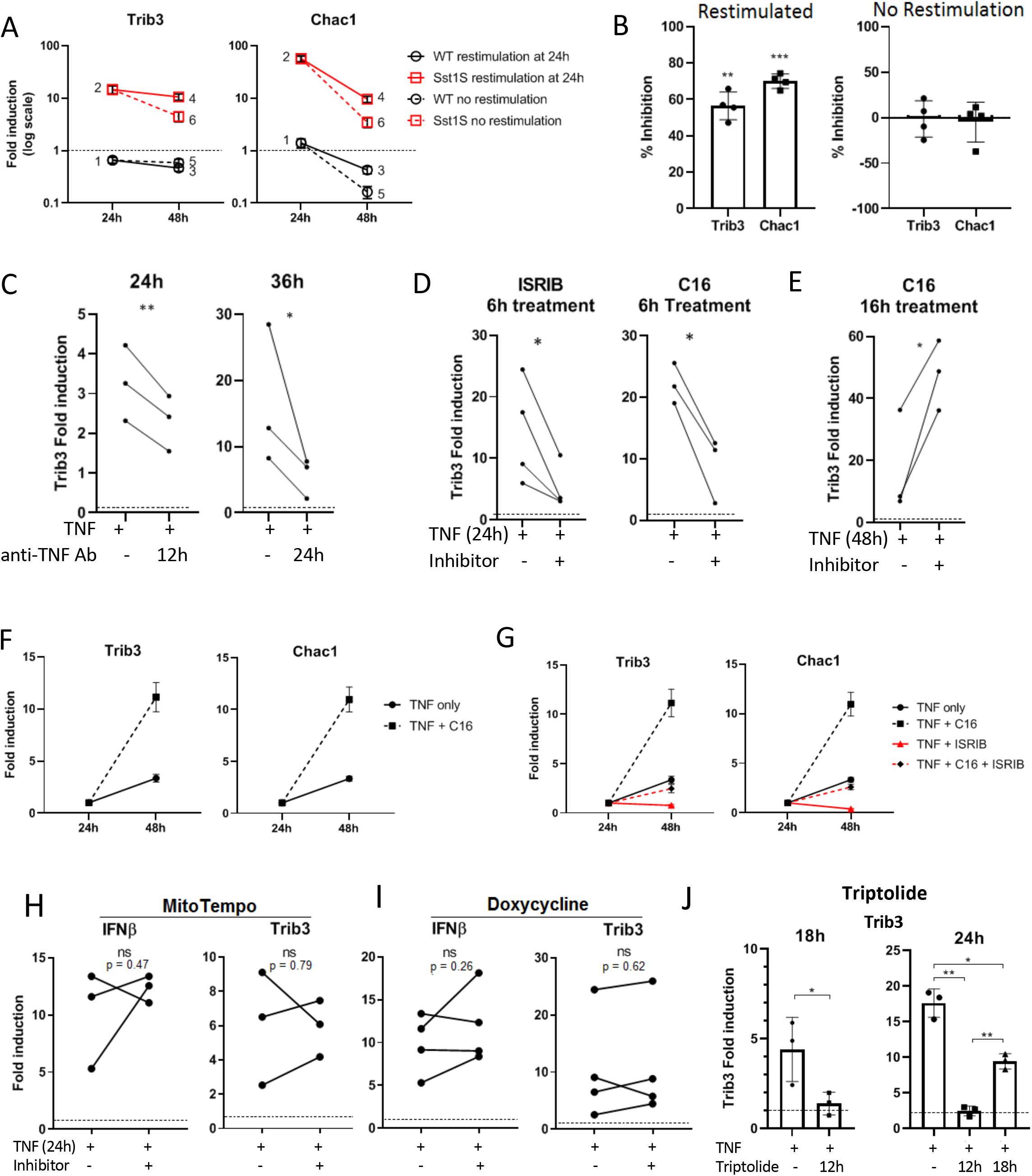
The Integrated Stress Response is activated in Sst1S, via PKR, in response to overarching, TNF-induced cellular stress. A. Sst1S and WT BMDMs were stimulated with 10 ng/mL TNF for 24 h, after which media was changed and cells were either re-stimulated with TNF or left unstimulated. Samples were harvested and qRT-PCR was performed after the first 24 h of stimulation, or 48 h after the original stimulation. See Figure 2A for experimental layout. One representative experiment shown of four performed, error bars represent SD of technical replicates. Significance data in supplemental table 1. B. Sst1S BMDMs were treated as in (A), except upon media change at 24 h, were also treated with anti-IFNAR antibody or isotype control. Fold induction between antibody and control were compared to determine percent inhibition. C. Sst1S BMDMs were stimulated with 10 ng/mL TNF for 24 or 36 h, or left untreated, with a TNF-blocking antibody added 12 h before harvest. qRT-PCR was performed on sample RNA to determine IFNb induction, compared to untreated samples (dotted line). Significance determined by paired t-test. D. Sst1S BMDMs were stimulated with TNF for 24 h. Inhibitors ISRIB (10 mM) or C16 (2 mM) were added 6 h before harvest (18 h of TNF stimulation) and analysis by qRT-PCR. Fold induction was calculated in the presence and absence of the inhibitors as a ratio to controls unstimulated with TNF. Significance determined by paired and ratio-paired t-test for C16 and ISRIB, respectively. E. Sst1S BMDMs were stimulated with TNF for 24h, then media was changed and BMDMs were restimulated with fresh TNF for an additional 24h (for a total of 48 h, as shown in Fig.2A). C16 was added 16h before harvest. One representative experiment shown, of two performed. Significance by paired t-test. F. Sst1S BMDMs were stimulated with TNF for 24 h, then media was changed and BMDMs were restimulated with fresh TNF as in E (for a total of 48 h before harvest). C16 (2uM) C16 was added during the whole re-stimulation period (24 h). qRT-PCR was performed for Trib3 and Chac1 to determine relative fold induction by TNF in the presence and absence of C16, as compared to their levels after 24 h of TNF stimulation. One representative experiment shown, of two performed. Significance by paired t-test. G. Sst1S BMDMs were stimulated with TNF for a total of 48h as in (E) and (F). The inhibitors (ISRIB at 10 mM and C16 at 2 mM) were added for the duration of the TNF re-stimulation (from 24 to 48 h) either separately or in combination. qRT-PCR was performed, and fold-induction was calculated as in (F). Single experiment shown, error bars represent standard deviation of technical replicates. Statistical analysis for all figure panels: *p < 0.05, ** p < 0.01, *** p < 0.001. H. Sst1S macrophages were treated with 10 ng/mL TNF or TNF plus MitoTempo ROS scavenger. qRT-PCR analysis was conducted for IFNβ and Trib3 and normalized to untreated controls. No significance observed for IFNβ or Trib3 by ratio-paired t tests. I. Sst1S macrophages were treated with 10 ng/mL TNF or TNF plus Doxycycline. qRT-PCR analysis was conducted for IFNβ and Trib3 and normalized to untreated controls. No significance observed for IFNβ or Trib3 by ratio-paired t-tests. J. Sst1S macrophages were treated with 10 ng/mL TNF for 18 or 24 h, or left untreated. Triptolide was added to indicated samples either 12 or 18 h after TNF. qRT-PCR analysis was conducted for Trib3 and normalized to untreated controls. Single experiment shown. Error bars represent standard deviation of technical replicates. Error bars and significance analysis based on technical replicates.

Indeed, inhibition of PKR reduced the expression of the ISR markers at 18 – 24 h of TNF stimulation (**Fig.3D**). Unexpectedly, PKR inhibition during the 24 – 48 h of TNF re-stimulation led to dramatic upregulation of Trb3 and Chac1 mRNA expression (**Fig.3E and 3F**), contrasting the IFNβ downregulation by the same treatment (**Suppl.Fig.2B**). This observation suggested to us that at this late stage the ISR and IFNβ regulation were uncoupled. To determine whether other pathways may be involved in the ISR increase in addition to PKR at this advanced stage of TNF response, we compared isolated and combined effects of C16 and ISRIB on ISR marker upregulation during the 24 – 48 h period. Unlike PKR inhibitor C16, ISRIB completely inhibited the TNF-induced ISR. Moreover, when combining C16 and ISRIB, we observed that ISRIB efficiently suppressed the C16-induced ISR escalation as well (**Fig.3G**). These observations suggested that during prolonged exposure to TNF and PKR blockade additional ISR pathways became activated.

We reasoned that the IFNβ ➢PKR ➢ISR axis must be activated in response to an ongoing stressor in TNF-stimulated Sst1S macrophages that emerges prior to PKR activation. In this scenario, PKR activation plays an adaptive role by reducing cap-dependent protein biosynthesis via eIF2α phosphorylation. Blocking PKR without resolving the underlying stressor, however, results in a consequent activation of alternative ISR pathways by other eIF2α kinases, such as PERK and HRI that are activated by misfolded proteins in ER and cytoplasm, respectively(26). Indeed, previously we observed an upregulation of heat shock protein genes Hspa1a and Hspa1b specifically in the Sst1S BMDMs, as early as 12 h of TNF stimulation pointing to the accumulation of misfolded proteins in cytoplasm prior to the ISR escalation. Treatment with ROS scavenger BHA prevented the protein aggregate accumulation and escalation of the Hsp1 expression (16).

#### Mitochondrial dysfunction during the course of TNF stimulation in Sst1S macrophages

Defective mitochondria could play a causative role in the aberrant Sst1S response to TNF via mitochondrial ROS production. Therefore, we compared mitochondrial function in quiescent and TNF-stimulated WT and Sst1S macrophages using standard Seahorse Extracellular Flux Analysis. We observed an increase in Basal Oxygen Consumption Rates (OCR) and ATP-linked respiration (ATP production) and reduced reserve capacity of mitochondria after 18 h of TNF stimulation in both WT and Sst1S macrophages, but no significant differences between the strains (**Suppl. Fig. 3A**). We did observe, however, a TNF-dependent strain difference in Extracellular Acidification Rate (ECAR) after inhibition of oxidative phosphorylation by FCCP and Antimycin. This difference, observable only in the absence of mitochondrial respiration, may reflect either a greater TNF-induced glycolytic rate in Sst1S cells or an impaired pH buffering capacity. Next, we excluded this hypothesis, because a mitochondria-targeted ROS scavenger MitoTempo (27) suppressed neither the IFNβ super-induction, nor the ISR marker expression (**Fig.3H**).

Another known mechanism linking mitochondrial damage to IFN-I and PKR activation is the leakage of mitochondrial RNA (mtRNA) to cytoplasm and activation of dsRNA sensors, including RIG-I and PKR (28). To test this hypothesis, we stimulated Sst1S macrophages with TNF in the presence of doxycycline, which is known to specifically inhibit mtRNA transcription. However, this treatment demonstrated no significant effects on either IFNβ or Trib3 induction by TNF (**Fig.3I**). In contrast, triptolide, an inhibitor of RNA polymerase II-mediated nuclear transcription (29), significantly reduced the Trib3 induction by TNF (**Fig.3J**). The triptolide effect was maximal when it was added at 12 - 18 h of TNF stimulation, i.e. during a critical interval of the aberrant IFNβ upregulation followed by the ISR escalation. Importantly, the triptolide treatment specifically inhibited the expression of ISR markers, as compared to IFNβ or housekeeping gene levels, thus excluding a possibility that its effect was non-specific due to a global transcription inhibition. The data also demonstrated that, at least initially, nuclear transcripts were recognized by PKR, but not other RNA sensors that, otherwise, would upregulate the IFNβ transcription in TBK1-dependent manner.

The above data demonstrate that mitochondrial damage does not initiate the aberrant response of the Sst1S macrophages to TNF. Nevertheless, after 48 h of TNF stimulation, we observed a morphological change of widened mitochondrial cristae in both WT and Sst1S with TNF (**Suppl.Fig.3B**). These morphological changes were more severe in Sst1S mitochondria, and were characterized by matrix “void” structures with absent cristae (**Suppl.Fig.3B**, bottom right). Taken together, our data demonstrate that greater mitochondrial damage caused by prolonged TNF stimulation in Sst1S macrophages is a consequence of, but not the cause, of their aberrant activation by TNF.

### Defective anti-oxidant response and free iron drive the aberrant response to TNF

#### The early hallmarks of the aberrant Sst1S macrophage response to TNF

To start revealing the deficiency that underlies the Sst1S phenotype, we compared global mRNA expression profiles of Sst1S and WT macrophages at 12 h of TNF treatment using RNA-seq (**Fig.4**). At this time point, we did not observe differential expression of ISR or IFN-I pathway genes between the Sst1S and WT cells. Gene set enrichment analysis (GSEA) of genes differentially expressed between the TNF-stimulated B6.Sst1S and B6 macrophages at this critical junction revealed that the mutant macrophages were deficient in detoxification of reactive oxygen species, cholesterol homeostasis, fatty acid metabolism and oxidative phosphorylation, while sumoylation and DNA repair related pathways were upregulated (**Supplemental Table 2**).

The candidate genes encoded within the *sst1* locus, Sp110 and Sp140, were strongly upregulated by TNF stimulation exclusively in the WT macrophages. In contrast, their expression in the B6.Sst1S BMDMs was severely reduced. Because both of those genes are putative regulators of chromatin organization and transcription (reviewed in (30)), we wanted to determine whether specific regulatory networks were associated with the candidate genes in TNF-stimulated macrophages. First, we inferred mouse macrophage gene regulatory network using the GENIE3 algorithm (31) and external gene expression data for mouse macrophages derived from Gene Expression Omnibus (GEO) (32). This network represents co-expression dependencies between transcription factors and their potential target genes, calculated based on mutual variation in expression level of gene pairs (33, 34). This network analysis revealed that Sp110 and Sp140 genes co-express with Nfe2l1 (Nuclear Factor Erythroid 2 Like 1, also known as Nrf1) and Mtf (metal-responsive transcription factor) TFs and their targets in mouse macrophages (**Fig. 4A**). Therefore, we postulated that the *sst1*-encoded Sp110 and/or Sp140 may be involved in the regulation of the Nfe2l1- and Mtf1-mediated pathways. These TFs are involved in regulating the response to oxidative damage and heavy metals, respectively. Thus, both GSEA and network analyses highlighted the potential differences in regulatory mechanism in response to oxidative stress in TNF-stimulated Sst1S and WT macrophages.

**Figure 4.**
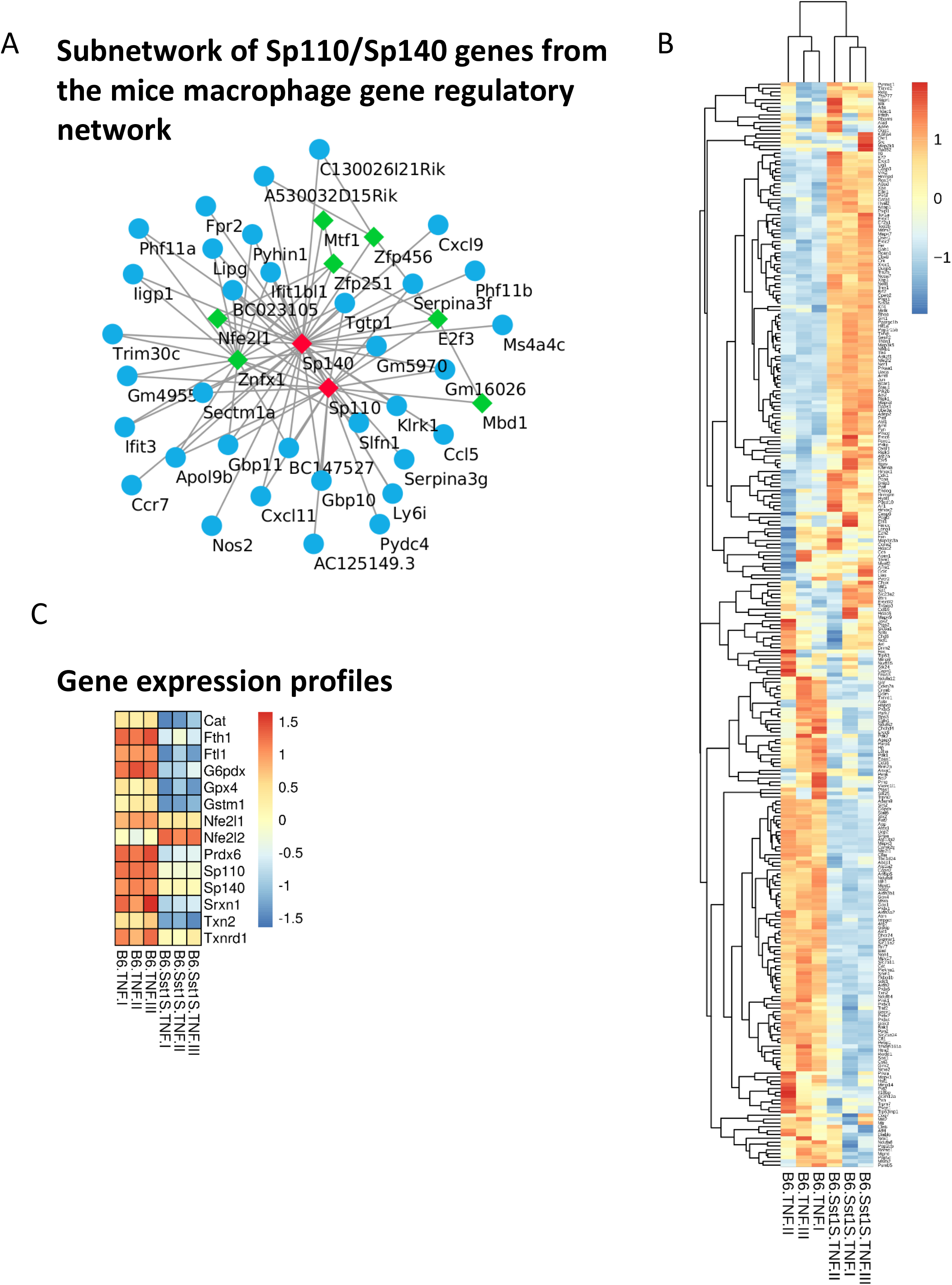
Gene expression profiling of Sst1S and WT BMDMs stimulated with TNF (10 ng/ml) for 12 h. A. Gene regulatory network analysis. The network represents a subnetwork of Sp110/Sp140 genes from the mice macrophage gene regulatory network. Only first neighbors of Sp110/Sp140 genes were selected. Green nodes represent transcription factors, blue nodes denote their potential targets. The mouse macrophage gene regulatory network was inferred using GENIE3 algorithm based on external gene expression data for mice macrophages derived from Gene Expression Omnibus (GEO). B. Analysis of genes related to oxidative stress. The heatmap was generated using FPKM values of genes related to response to oxidative stress (gene ontology category GO: 0006979). C. Gene expression profiles for a list of the *sst1* controlled genes. The heatmap was generated using RNA-seq expression profiles of Sst1S and WT macrophages at 12 h of TNF treatment. For heatmap generation, FPKM values were scaled using Z-scores for each tested gene.

To further test this hypothesis, we analyzed the expression of a gene ontology set “response to oxidative stress” (GO0006979, 416 genes, **Fig.4B**). We observed clear separation of these genes in two *sst1*-dependent clusters. Many known anti-oxidant genes were upregulated in TNF-stimulated WT macrophages to a significantly higher degree as compared to the Sst1S mutants. Functional pathway profiling revealed that B6-specific cluster represents genes involved in “detoxification of reactive oxygen species”, “Ferroptosis”, “HIF-1” and “peroxisome” signaling pathways, while Sst1S-specific cluster contains genes from “MyD88-independent TLR4”, “DNA repair”, “Oxidative Stress Induced Senescence” and “SUMOylation” pathways (**Suppl. Table 3**). Transcription factor binding site analysis of genes from the B6-specific cluster revealed an enrichment of Nfe2l1/Nfe2l2 binding site sequence motifs. In contrast, overrepresentation of E2F, Egr1 and Pbx3 transcription factor binding sites was found for genes from Sst1S-specific cluster (**Suppl. Table 4).** A master regulator analysis also revealed a key role for Nfe2l (NF-E2-like) transcription factors (TFs) as regulators of genes differentially induced by TNF in WT and Sst1S BMDMs (**Suppl. Table 5**). However, in the B6 phenotype Nfe2l1 was upregulated to a greater extent, as compared to Nfe2l2, while a reverse relationship was observed in the mutant (**Fig. 4C**). This difference may explain preferential utilization of the Nfe2l TFs in B6 and Sst1S. The Nfe2l1 and Nfe2l2 TF are known to bind to promoters of an overlapping, but not identical, set of genes that contain anti-oxidant response elements (ARE) in their promoters (35). Their upstream regulators, however, are different(36). Thus, the superior B6 response to TNF-induced oxidative stress may be driven by preferential utilization of the Nfe2l1- and Mtf-mediated pathways.

#### Antioxidant blockade and iron chelation correct the aberrant macrophage activation

Because oxidative damage is a well-established inducer of JNK (37), we hypothesized that the JNK-mediated hyperactivation of the IFN-I and ISR in TNF-stimulated Sst1S macrophages was driven by oxidative stress. Indeed, treatment of Sst1S macrophages with the anti-oxidant compound BHA during TNF stimulation reduced the IFNβ and Trib3 transcript levels to almost background levels (**Fig.5A**). Furthermore, the increased cell death rates induced by 50 ng/mL of TNF could also be reversed with BHA (**Fig.5B**).

**Figure 5:**
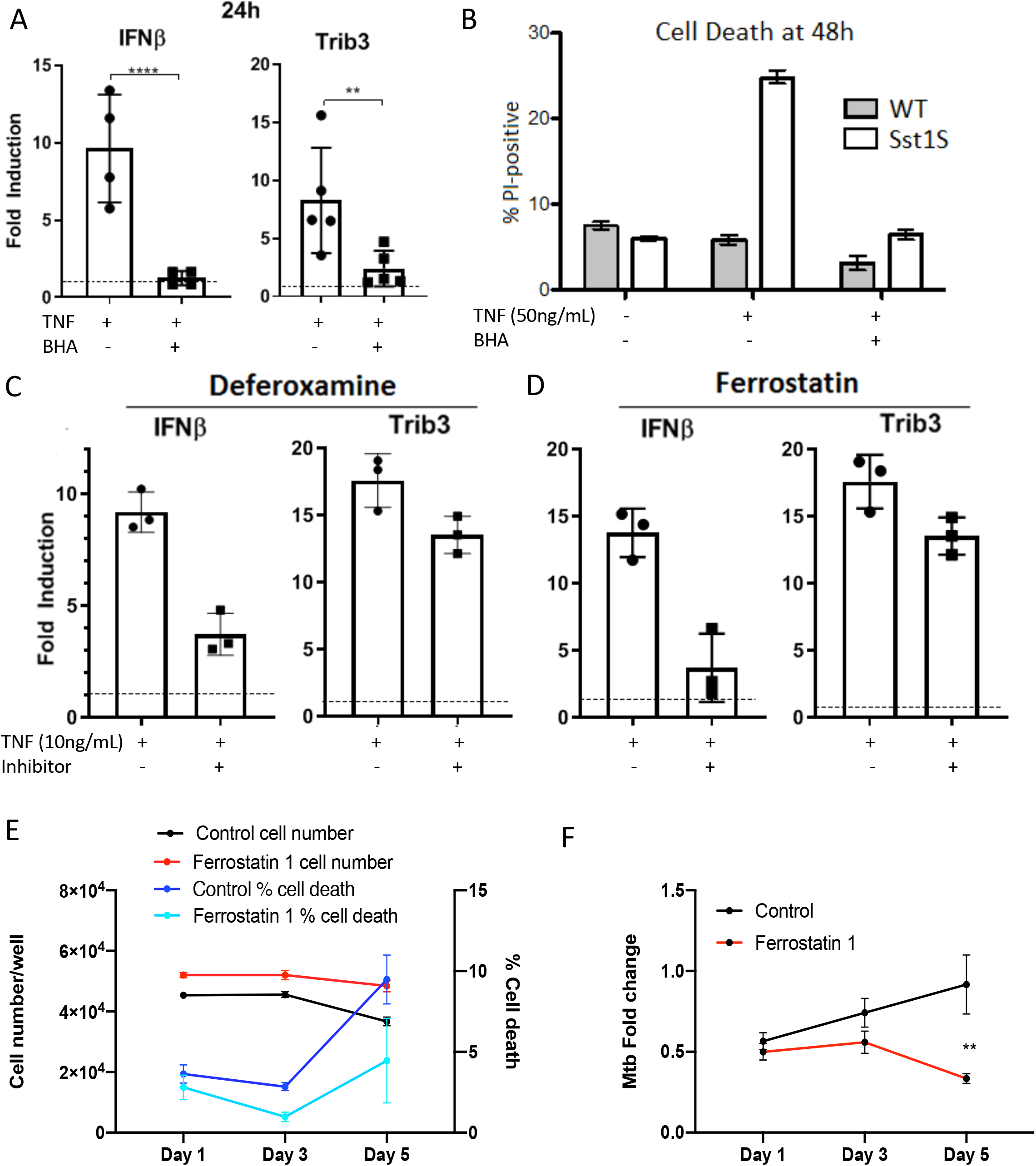
The Sst1S phenotype and susceptibility to Mtb is dependent upon iron-induced oxidative stress. A. BHA pretreatment for 24 h before and throughout 24 h of TNF stimulation in Sst1S BMDMs reduces IFN and Trib3 RNA induction, relative to untreated controls. Significance by ratio-paired t-test. B. Sst1S and WT were treated for 48 h with 50 ng/mL TNF, TNF plus BHA, or left untreated, and analyzed by PI staining and fluorescent cell counting to determine cell death rates. Single representative experiment shown, of three repeat experiments. Error bars represent standard deviation of technical replicates. C. Sst1S BMDMs were treated with TNF or TNF plus deferoxamine for 24 h, and compared to untreated controls by qRT-PCR for IFNβ and Trib3. Single representative experiment shown, additional experimental data in supplemental tables 7 and 8. Error bars represent standard deviation of technical replicates. D. Sst1S BMDMs were treated with TNF or TNF plus Ferrostatin for 24 h, and compared to untreated controls by qRT-PCR for IFNβ and Trib3. Single representative experiment shown, additional experimental data in supplemental tables 9 and 10. Error bars represent standard deviation of technical replicates. E. B6.Sst1S macrophages were pretreated with TNFα (10 ng/mL) for 8 h and subsequently treated with TNFα in combination with Ferrostatin 1 (3 μM) for 16 h. The cells were infected with Mtb at MOI 0.5 and harvested at 1, 3 and 5 days post-infection and analyzed for cell survival and death. Error bars indicate standard deviation of technical replicates, significance values determined by 2-way ANOVA with multiple comparisons. F. B6.Sst1S macrophages were treated as in (E) and harvested at 1, 3 and 5 days post-infection. The Mtb load was determined using a qPCR-based method, normalized to a BCG spike for internal control. Error bars indicate standard deviation of technical replicates, significance values determined by 2-way ANOVA with multiple comparisons.

We noted that ferritin light (Ftl) and heavy (Fth) chains were among the most abundant transcripts upregulated by TNF at 12 h in WT macrophages. However, TNF stimulation had opposite effects on the Ftl and Fth gene expression in B6.Sst1S cells: both genes were downregulated at 12 h. Because free intracellular iron has potent catalytic activity to produce damaging hydroxyl radicals from peroxide via Fenton reaction(38), we hypothesized that ROS produced in TNF-activated macrophages may lead to greater damage in the Sst1S background. We tested this hypothesis by adding deferoxamine, an iron chelator, to Sst1S macrophages along with the standard TNF treatment for 24 h. Indeed, we observed a decrease in both IFNβ and Trib3 in iron-chelated samples compared to TNF alone (**Fig.5C)**. Because defects in iron metabolism in macrophages lead to accumulation of lipid peroxidation products and eventually cell death by ferroptosis(39), we also tested ferrostatin 1, an inhibitor of lipid peroxidation in the ferroptosis pathway(40). Ferrostatin treatment along with TNF for 24 h also reduced IFNβ and Trib3 levels compared to TNF alone (**Fig.5D**). We also confirmed the beneficial effects of ferrostatin on Mtb-infected B6.Sst1S macrophages - increasing macrophage cell survival (**Fig.5E**) and limiting the Mtb growth (**Fig.5F**). Ferrostatin produced the greatest improvement in infection outcomes over 5 days of infection at a concentration of 3 mg/mL, but was also effective at lower doses (**Suppl.Fig.4**). These data demonstrate that failure to upregulate pathways known to prevent lipid peroxidation and ferroptosis (ferritin and Gpx4) by Sst1S macrophages in response to activation with TNF predisposes them to abnormal IFN-I pathway activation and, likely, accumulation of oxidative damage products to sublethal levels. While not inducing macrophage death per se, this ultimately leads to poorer outcomes during Mtb infection. A unified model of cascades underlying the aberrant response of the Sst1S macrophages to TNF is depicted in Fig.6.

**Figure 6.**
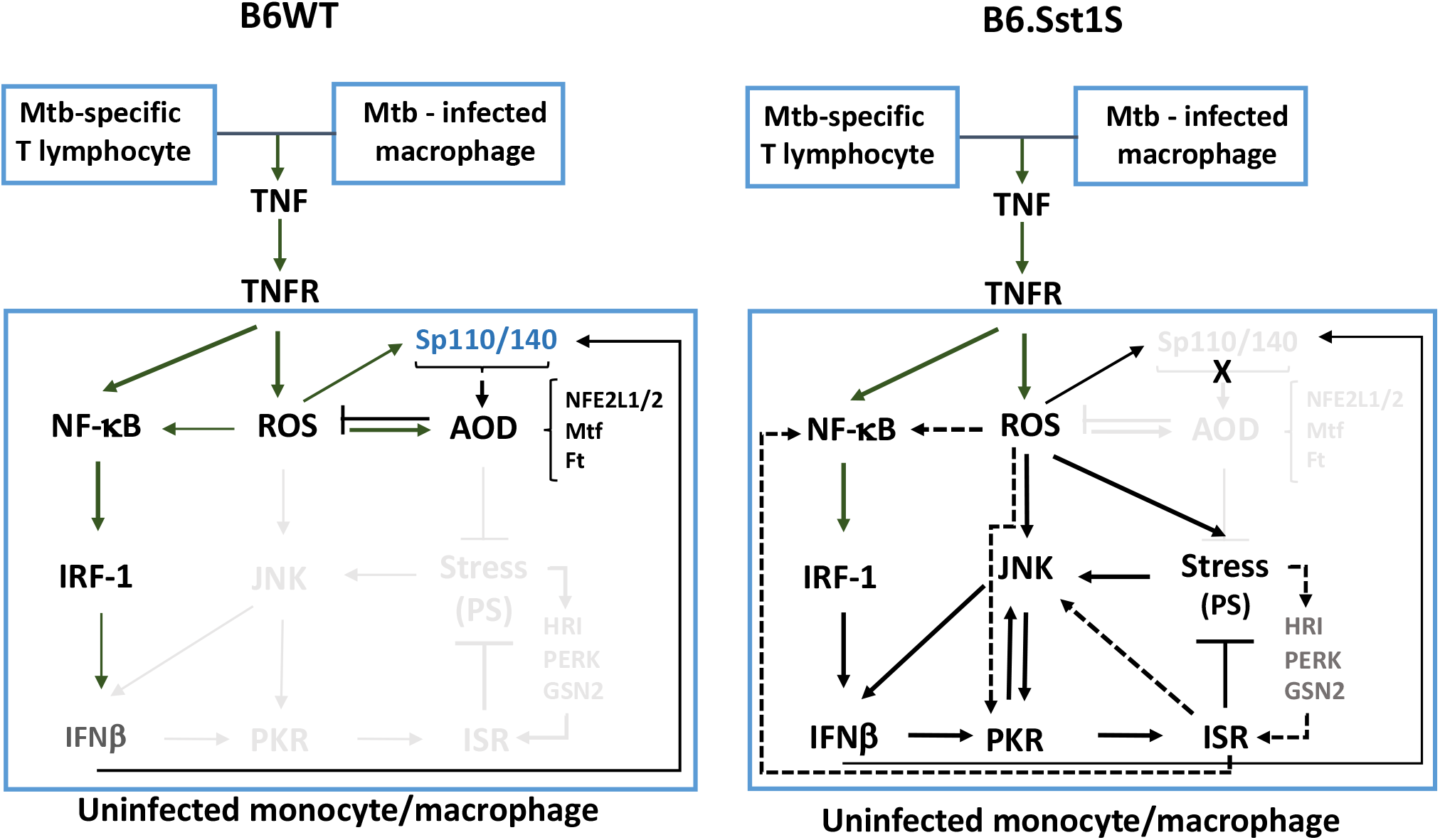
Adaptive (B6wt) and maladaptive (B6.Sst1S) circuits of TNF response. A. The B6wt macrophages display the upregulation of anti-oxidant defense pathways (AOD) in the *sst1*-dependent manner and a modest upregulation of IFNb mediated by IRF1. B. The B6.Sst1S mutant macrophages fail to activate adequate AOD, experience unresolving oxidative stress, including proteotoxic stress (PS), and upregulate a stress-activated JNK pathway. This results in the IFNb super-induction, PKR activation and the development of the integrated stress response (ISR), as an alternative PS mitigation strategy. The above cascades lead to the formation of positive feedback loops sustaining the unresolving stress and the type I IFN pathway hyperactivation (solid lines). Dashed lines depict additional feedforward and positive feedback mechanisms based on current literature.

## DISCUSSION

### Iron-driven ROS is a driver of Mycobacterial susceptibility and Type I Interferon pathway hyperactivity

Macrophages are the main cell population responsible for the *sst1*-mediated phenotype in vivo and in vitro (13, 20, 21). Sst1-susceptible macrophages exhibit an atypical response to prolonged TNF stimulation characterized by an upregulation of the IFN-I pathway driving the ISR via PKR activation(16). Here we demonstrate that upstream of IFNβ superinduction there occurs a critical dysregulation of a free iron-dependent oxidative stress, in which a failure of the antioxidant response and iron control mechanisms results in unresolving stress in Sst1S macrophages. We observed gradual escalation of the dysregulated TNF response dominated by a hyperactive IFN-I pathway. The later was inhibited by ferrostatin-1, a ferroptosis inhibitor that specifically inhibits lipid peroxide formation (40). Removing excess of free iron using a synthetic iron chelator deferxoamine (DFOM) produced similar effects. These data suggest that benefit we observe with DFOM and ferrostatin-1 use in Mtb-infected Sst1S macrophages is not inhibition of ferroptosis, but prevention of the damaging lipid peroxidation cascade. Recently DFOM treatment was shown to enhance glycolysis and enhance immune activation of primary human macrophages infected with virulent Mtb in vitro(41). Ferrostatin-1 was used in vivo to specifically target necrosis in mouse TB lesions(11).

Of note, downregulation of Fth light and heavy chains and ferroptosis inhibitors in TNF-stimulated Sst1S macrophages occurs in a coordinated manner, thus creating an intracellular environment for increased generation of free radicals and their unopposed spread. Likely, this potentially suicidal program may enhance non-specific free radical attack on intracellular pathogens. However, the survival of Mtb does not appear to be negatively affected by an excess of iron and peroxide products due to protection by mycobacterial cell wall and multiple mycobacterial anti-oxidant pathways which promote mycobacterial survival even in highly oxidative environments (42, 43). In contrast, free radicals damage the mutant macrophages. After the removal of damaging lipid peroxides by ferrostatin treatment, we observed increased macrophage survival and reduced mycobacterial proliferation. Thus, protecting TNF-stimulated Sst1S macrophages from self-inflicted oxidative damage enables them to continue as functional effectors of innate immunity, rather than merely persisting as fodder for the growing infection.

Our findings demonstrate that the *sst1* locus plays a major role in coordinating an anti-oxidant response with macrophage activation. Notably, expression of Fth light and heavy chains, catalase, selenoprotein genes thioredoxin reductase (Txnrd1) and glutathione peroxidases (Gpx1 and Gpx4) are coordinately upregulated after TNF stimulation in the *sst1*-dependent manner. The glutathione peroxidase 4 (GPX4) protein plays a central role in preventing ferroptosis because of its unique ability to reduce hydroperoxide in complex membrane phospholipids and, thus, limit self-catalytic lipid peroxidation leading to ferroptosis(39). Thus, the wild type allele of the *sst1* locus encodes positive regulators of the ferroptotic pathway inhibitors whose upregulation increase macrophage stress resilience. Recent in vivo studies demonstrated that ferritin and an anti-oxidant pathways were upregulated in inflammatory macrophages isolated from Mtb-infected B6wt mice(44). Inactivation of these regulators, as exemplified by the mutant Sst1S phenotype, promotes oxidative damage by coordinately increasing free iron pool and non-enzymatic free radical production, and simultaneously decreasing buffering capacity of intracellular antioxidants. Therefore, we propose that the *sst1* locus encodes factors that, in certain contexts, are critical for the macrophage fate determination.

### A mechanism to connect anti-oxidant response dysregulation to the sst1 gene locus

A molecular mechanism of the anti-oxidant response regulation by the *sst1* locus in response to TNF remains an unanswered question. However, this study provides new evidence supporting a major role of the Sp100 family genes encoded within the *sst1* locus in this phenotype. The Sp110 and Sp140 genes are known as interferon-inducible genes. Both genes, however, are not upregulated in response to TNF, IFNβ and IFNγ in macrophages isolated from mice that carry the *sst1* susceptibility allele, as opposed to the wild type.

Overexpression of Sp110 in the *sst1* susceptible macrophages and mice increased their resistance to Mtb(13). More recently, using CRISPR knockout, Vance and colleagues have shown that the Sp140 gene may play a bigger role in the *sst1*-mediated phenotype in mice(14). The human homologs to both of these genes are active in immune cells including macrophages (30, 45). Remarkably, genome wide association studies in human populations found that Sp140 polymorphisms were associated with two major chronic inflammatory diseases, Crohn’s Disease and Multiple Sclerosis (reviewed in (30). Human SP110 and SP140 are also a known Interferon-stimulated gene. Indeed, meta-analysis indicates that the expression of SP110 and SP140 is up-regulated in many disorders characterized by increased type I IFN activity (see supplemental Table 4).

Both Sp110 and Sp140 are nuclear chromatin-binding proteins with complex domain structures including histone- and DNA-binding domains. The SP140 protein contains a bromodomain, and has been observed to localize to H3K4me3- and H3K27me3-rich promoter and enhancer sites. These data and gene expression patterns indicate that Sp140 acts as transcriptional repressor in LPS stimulated macrophages (30, 46). Similarly, we have determined that mouse Sp110b acts as a non-specific transcriptional repressor(47).

To approach mechanisms of the Sp110 and Sp140-mediated transcriptional regulation, we constructed the Sp110 and Sp140 co-regulated gene networks using public databases and analyzed differentially expressed genes between the TNF-stimulated WT and Sst1S macrophages at 12 h, a critical divergence point between the two phenotypes. These analyses revealed a key role for NF-E2-like transcription factor Nfe2l1 as a regulator of genes upregulated by TNF in *sst1* resistant macrophages. Interestingly, in the wild type phenotype the Nfe2l1 mRNA was upregulated to a greater extent, as compared to Nfe2l2. The Nfe2l1 and Nfe2l2 (Nrf2) TFs are known to bind to promoters of an overlapping, but not identical, set of genes that contain anti-oxidant response elements (ARE) in their promoters (35). Their upstream regulators, however, are different(36). Of note, expression of a classical Nrf2 target gene heme oxygenase 1 were similar in both backgrounds. Thus, the superior anti-oxidant response of the *sst1* resistant macrophages may be driven by preferential utilization of the Nfe2l1-mediated pathways.

In contrast, promoters of the genes upregulated by TNF in the Sst1S background are enriched in the binding sites of pro-oncogenic homeobox domain transcription factor PBX3, and transcription factors involved in cell cycle progression and response to growth factors E2F and Egr1. Ferritin expression is known to be downregulated by oncogenes and *c-myc* (48). Thus, the WT and Sst1S macrophages display antagonistic programs – an anti-oxidant response or a cell growth, respectively. We propose that the *sst1*-encoded Sp110 and/or Sp140 family proteins regulate these choices in a context-dependent manner to fine-tune macrophage transcriptional responses for optimal stress adaptation or free radical production and self-destruction.

### Dual role of JNK - IFN – PKR circuit in the aberrant TNF response

Previously we demonstrated that IFNβ superinduction after 12 h of TNF stimulation in the Sst1S macrophages was mediated by synergy of NF-kB- and JNK-mediated pathways(16). Our new data traces the origin of the IFN-I hyperactivity to chronic oxidative stress, as it has been shown to activate ASK1 – JNK axis in many studies(49). Unexpectedly, we found no evidence of STING and TBK1-mediated pathway involvement including the in vivo observation that inactivation of those pathways had no effect on TB susceptibility of the B6.Sst1S mice(16, 17, 50).

As we explored the propagating mechanisms of the TNF response dysregulation beyond 16 h, we confirmed that TNF and JNK remain primary drivers of IFNβ at both early and late time points, but also discovered a more complex role for PKR. PKR is known to be a mediator of Integrated Stress Response activation by IFNβ in Sst1S, but it was not known to reciprocally upregulate IFNβ. We excluded ISR-mediated feedback as a primary mechanism due to the observation that global ISR inhibition by ISRIB did not affect IFNβ levels. Furthermore, inhibiting either JNK or PKR after 24 h completely abolishes IFNβ superinduction, and there is no additive effect because inhibition of either results in near 100% reduction of IFNβ levels relative to baseline. This supports the hypothesis that PKR and JNK are mechanistically in sequence and suggests the PKR - JNK connection(51) as a feasible mechanism for PKR feedback to IFNβ. Since a TNF blockade also abolishes IFNβ induction, we proposed that TNF, JNK and PKR are all in the same mechanistic pathway as follows: TNF in combination with JNK, activated by unopposed TNF-induced ROS, cause the IFNβ superinduction seen after 12 h. High levels of IFNβ induce PKR, which becomes activated and signals back to maintain JNK activation thus establishing a feedforward cycle maintaining the IFN-I hyperactivity and chronic ISR.

We do not yet know the source of PKR activation in our model. Recently it has been shown that PKR may be activated under cell intrinsic stresses by endogenous RNAs, including mitochondrial dsRNA and structured nuclear ssRNA, endoretroviral RNAs (52, 53), and snoRNAs (54). In addition, there is a known protein activator of PKR (PACT) that functions in an RNA-independent manner (55, 56). PACT is known to activate PKR in response a variety of cellular stresses (57), including ER stress (58) and oxidative stress (59, 60). Thus, the JNK – IFN – PKR circuit may serve as an integrator of various extrinsic and intrinsic stress signals.

In our model, it is activated via cell intrinsic pathway in response to unresolved oxidative stress in a step-wise manner. Initially, PKR expression is induced in response to IFNβ super-induction. This sensitizes the PKR branch of ISR to subsequent endogenous activating stimuli (RNA or PACT). At this stage, PKR is activated earlier than other eIF2α kinases and plays an adaptive role by reducing overall protein translation and, thus, preventing a more severe proteotoxic stress. In this scenario, the JNK – IFNβ - PKR – ISR branch represents a backup mechanism activated when an earlier anti-oxidant response fails. Eventually, however, it establishes and maintains a vicious cycle of unresolving ISR via PKR – JNK – IFN feedback loop. The chronic IFN-I pathway hyperactivity produces immunosuppressive and damaging effects at cellular (macrophage), tissue (granuloma) and systemic levels(17, 61–64).

### Do mechanisms underlying necrosis in TB granulomas controlled by the sst1 locus in mice provide generalizable insights into TB pathogenesis in humans?

In regard to specific mechanisms of human TB pathogenesis, our findings demonstrate that defect in macrophage response to TNF may underlie slow progression of chronic TB granulomas towards the necrotization in immunocompetent hosts. A central role of TNF in host resistance to TB and especially in organization and maintenance of TB granulomas is well established in all animal species(9, 65). However, TNF is often compared to a double-edged sword, because its excess is detrimental to the host and promotes progression towards disseminated TB disease(66–68). Our study links the development of macronecrotic TB granulomas in vivo with a more subtle differences in macrophage responses to TNF: in susceptible hosts, TNF even at relatively low levels may be sufficient to trigger a cascade of sublethal unresolving stress that can exacerbate the effects of additional stressors or infection.

Persistent TNF stimulation is required to maintain the full effects of dysregulation in Sst1S, and this constant stimulation model in vitro mimics the granuloma environment in vivo. Activated macrophages inside an established granuloma produce a supply of TNF for newcomer macrophages, which encounter the TNF on the granuloma periphery (69), and can become dysregulated before ever encountering the mycobacteria. Our studies suggest that innate or acquired dysregulation of an anti-oxidant defense does not lead to overt systemic immune deficiency. Instead, as granulomas gradually progress, macrophage crowding and exclusion of T lymphocytes from the inner granuloma core ensure that cell autonomous macrophage defects start playing more prominent roles leading to granuloma expansion and progression towards macronecrosis and cavities.

Similar macrophage dysregulation is likely to be present within a spectrum of TB disease in its natural human hosts and play an important role in the development of highly transmissible pulmonary TB in immunocompetent humans. From the pathogen’s evolutionary perspective, this trajectory would allow for the establishment of controlled chronic infection but prevent the progression toward systemic lethal disease for a period of time sufficient to allow for Mtb transfer to multiple individuals. Therefore, the Sst1S mouse model of necrotic TB granulomas can be used to further our understanding of Mtb virulence mechanisms, as well as to develop and test novel host-directed therapies specifically targeting the host and the pathogen pathways driving granuloma necrotization.

## MATERIAL AND METHODS

### Reagents

Recombinant mouse TNF was purchased from Peprotech. Recombinant mouse IL-3 was purchased from R&D. Mouse monoclonal antibody to mouse TNF (MAb; clone XT22), Isotype control and mouse monoclonal antibody to mouse IFNβ (Clone: MAR1-5A3) were purchased from Thermo scientific. SP600125 and C-16 purchased from Calbiochem. PKR inhibitor structural control (CAS # 852547-30-9) purchased from Santa Cruz Biotechnology. BHA obtained from Sigma. FBS for cell culture medium obtained from HyClone. Middlebrook 7H9 mycobacterial growth medium and 7H10 agar plates were made in house according to established protocols.

### Animals

C57BL/6J inbred mice were obtained from the Jackson Laboratory (Bar Harbor, Maine, USA). The B6J.C3-*Sst1*^*C3HeB/Fej*^ Krmn congenic mice were created by transferring the *sst1* susceptible allele from C3HeB/FeJ mouse strain on the B6 (C57BL/6J) genetic background using twelve backcrosses (referred to as B6.Sst1S in the text). All experiments were performed with the full knowledge and approval of the Standing Committee on Animals at Boston University in accordance with relevant guidelines and regulations.

### Tissue Inactivation, Processing, and Histopathologic Interpretation

Tissue samples were fixed for 72 h in 4% paraformaldehyde, at which time tissues were removed from the BSL-3 laboratory and stored in 1X PBS at 4°C until processed in a Leica PELORIS automated vacuum infiltration processor (Leica, Wetzlar, Germany), followed by paraffin embedding with a Histocore Arcadia paraffin embedding machine (Leica). Paraformaldehyde-fixed, paraffin-embedded blocks were sectioned to 5 μm, transferred to positively charged slides, deparaffinized in xylene, and dehydrated in graded ethanol. Tissues sections were stained with hematoxylin and eosin (H&E) for histopathology and a New Fuchsin Method to detect Acid-Fast Bacilli (Poly Scientific R&D Corp Catalog #K093-16OZ, Bay Shore, NY).

### Chromogenic Monoplex Immunohistochemistry

A rabbit specific HRP/DAB detection kit was employed (Abcam catalog #ab64261, Cambridge, United Kingdom). In brief, slides were deparaffinized and rehydrated, endogenous peroxidases were blocked with hydrogen peroxidase, antigen retrieval was performed with a citrate buffer for 40 minutes at 90°C using a NxGen Decloaking chamber (Biocare Medical, Pacheco, California), non-specific binding was blocked using a kit protein block, the primary antibody was applied at a 1.200 dilution, which cross-reacts with mycobacterium species (Biocare Medical catalog#CP140A,C) and was incubated for 1 h at room temperature, followed by an anti-rabbit antibody, DAB chromogen, and hematoxylin counterstain. Uninfected mouse lung was examined in parallel under identical conditions with no immunolabeling observed serving as a negative control.

### BMDMs culture and Treatment

Isolation of mouse bone marrow and culture of BMDMs were carried out as previously described (13). TNF-activated macrophages were obtained by culture of cells for various times with recombinant mouse TNF (10 ng/ml).

BMDM were seeded in tissue culture plates. Cells were treated with TNF and incubated for 24 h at 37°C with 5% CO2. The media were replaced with new media and the cells were either incubated with TNF (Re-stimulation) or without TNF for additional 24 h at 37°C with 5% CO2. PKR inhibitor (C-16) 2 μM final, ISR inhibitor (ISRIB) 10 mM final, a-IFNAR and a-Isotype antibody (1:100) were added 24 h of TNF treatment with the media replacement.

### Infection with *M. tuberculosis*

For infection experiments, M. tuberculosis H37Rv was grown in 7H9 liquid media for three days, harvested, and diluted in media without antibiotics to the appropriate MOI. 100 μL of M. tuberculosis-containing media at appropriate MOIs were added to BMDMs grown in a 96-well plate format and pre-treated for 24 h with TNF, if applicable. The plates were then centrifuged at 500g for 5 minutes and incubated for one h at 37 o C. Cells were then treated with Amikacin at 200 μg/μL for 45 minutes, to kill any extracellular bacteria. Cells were then washed and cultured with TNF and inhibitors as applicable in DMEM/F12 containing 10% FBS medium without antibiotics at 37 ° C in 5% CO 2 for the period described in each experiment, with media change and replacement of TNF and/or inhibitors every 48 h. MOI were confirmed by colony counting on 7H10 agar plates. All procedures involving live *M. tuberculosis* were completed in Biosafety Level 3 containment, in compliance with regulations from the Environmental Health and Safety at the National Emerging Infectious Disease Laboratories, the Boston Public Health Commission, and the Center for Disease Control.

### Cytotoxicity and Mycobacterial growth assays

Isolated and treated BMDMs were stained with Hoechst or Propidium Iodide nuclear staining. Media was removed and the staining reagent was added to the cells at a 1:100 dilution in PBS for 30 minutes, after which cells are washed with PBS and imaged directly using the Celigo microplate cytometer (Nexcelom). For viability tests in infected cells, macrophages were cultured in 96-well plates and infected with Mtb as described above. At harvest samples were treated with Live-or-Dye™ Fixable Viability Stain (Biotium) at a 1:1000 dilution in PBS/1% FBS for 30 minutes. After the stain, samples were gently washed to ensure no loss of dead cells from the plate, and fixed with 4% Paraformaldehyde for 30 minutes. The fixative was again washed and replaced with PBS, after which sample plates were decontaminated for removal from containment. Images and cell counts for both infected and uninfected cells were acquired using a Celigo microplate cytometer (Nexcelom). The intracellular bacterial load was determined by quantitative real time PCR (qPCR) using specific set of Mtb and *M.bovis*-BCG primer/probes with BCG spikes added as internal control (details of the method has described in Yabaji et al., submitted for publication).

### RNA Isolation and quantitative PCR

Total RNA was isolated using the RNeasy Plus mini kit (Qiagen). cDNA synthesis was performed using the High-Capacity cDNA Reverse Transcription Kit (ThermoFisher), or was completed during one-step qPCR using SuperScript II Reverse Transcriptase (Invitrogen). Real-time PCR was performed with the GoTaq qPCR Mastermix (Promega) using the CFX-96 real-time PCR System (Bio-Rad). Oligonucleotide primers were designed using Primer 3 software (supplemental table 13) and specificity was confirmed by Primer efficiency analysis and melting curve analysis. Thermal cycling parameters involved 40 cycles under the following conditions: 95 °C for 2 minutes, 95 °C for 15 s and 60 °C for 60 s. Each sample was set up in triplicate and normalized to 18S expression by the ddCt method.

### Analysis of RNA sequencing data

Raw sequence reads were mapped to the reference mouse genome build 38 (GRCm38) by STAR(70). The read count per gene for analysis was generated by featureCounts (71). Read counts were normalized to the number of fragments per kilobase of transcript per million mapped reads (FPKM) using the DESeq2 Bioconductor R package(72). Pathway analysis was performed using the GSEA method implemented in the camera function from the limma R package(73). Databases KEGG, MSigDB Hallmark, Reactome were used for the GSEA analysis. Transcripts expressed in either 3 conditions with FPKM > 1 were included in the pathway analyses. To infer mice macrophage gene regulatory network, we used ARCHS4 collected list of RNE-seq data for mice macrophages cells from Gene Expression Omnibus (GEO)(74). The total number of analyzed experimental datasets was 1960. This gene expression matrix was utilized as input for the GENIE3 algorithm, which calculated the most likely co-expression partners for transcription factors. The list of transcription factors was derived from the Animal TFDB 3.0 database(75). The Virtual Inference of Protein Activity by Enriched Regulon Analysis (VIPER) algorithm was used to estimate the prioritized list of transcription factors(76). The software Cytoscape (version 3.4.0) was used for network visualization(77). The i-cisTarget web service was used to identify transcription factor binding motifs that were over-represented on a gene list(78). Statistical analysis and data processing were performed with R version 3.6.1 (https://www.r-project.org/) and RStudio version 1.1.383 (https://www.rstudio.com).

### Western Blot Analysis

SDS-PAGE was completed on 8% Acrylamide Bis-Tris gels, self-made according to standard protocols. Whole cell lysates were prepared using RIPA lysis buffer (Thermo Scientific) with added protease and phosphatase inhibitors. Gels were run with 50ug total protein per well, as determined by BCA assay, and transferred to PVDF membrane (Millipore). After blocking for 2 h in blocking buffer (Li-Cor) or 5% skim milk in TBS-T buffer [20 mM Tris-HCl (pH 7.5), 150 mM NaCl, and 0.1% Tween20]. the membranes were incubated overnight with primary antibody at 4° C. Bands were detected by chemiluminescence (ECL) kit (Thermo Scientific) or by Li-Cor Odyssey for fluorescent antibodies. Loading control β-actin (1:2000, Sigma) was evaluated on the same membrane. PKR antibody (Santa Cruz) was used at a dilution of 1:200, and phospho-PKR (Thr 446) antibody (Santa Cruz) was used at a dilution of 1:150. Secondary antibodies used include fluorescent goat anti-mouse (1:20,000) and anti-rabbit (1:20,000) (Li-Cor), and HRP-conjugated goat anti-mouse (1:8000) and anti-rabbit (1:8000).

### Seahorse Extracellular Flux Analysis

BMDMs were plated on Seahorse-specific 96-well culture dishes (obtained from Agilent) at a density of 12,000 cells in 200uL of DMEM/F12. Culture conditions were as described above. Seahorse analysis was completed as per the standard Agilent Stress Test Protocol 43 with standardized Oligomyin and Antimycin, and an FCCP concentration of 2uM.

### Transmission Electron Microscopy

Macrophages were placed in 1x PBS on ice for 5 minutes to loosen adhesion to cell plate, then scraped into solution. Cells were pelleted at 300g for 5 minutes at RT, then fixed in a solution of 4% PFA. Fixed cells were collected by centrifugation and resuspended in a low-melting point agarose solution at 42°C, then pelleted and allowed to cool and solidify. Fixed cell pellets in agarose gel were then post-fixed with 1% osmium tetroxide in 0.15M cacodylate buffer, dehydrated in 50, 90 and 100% serial acetone, and embedded in epoxy resin. Semi-thin sections were cut at 2 mm with a glass knife, and stained with toluidine blue for light microscopic examination. After a suitable area was chosen, a diamond knife was used to cut ultrathin sections, about 72 nm, which were then mounted on 200-mesh cleaning copper grids and stained with 4% uranyl acetate and 0.4% lead citrate. The sections were examined in a JEOL JEM-1010 transmission electron microscope and different magnification images were obtained with an Erlangshen ES100W digital camera (Gatan, Pleasanton, CA).

### Statistical Analyses Overview

Cell Survival, Cell death, qPCR data, protein quantification data and composite Seahorse data were analyzed by paired t-tests, or one- or two-way ANOVA with post hoc tests for multiple comparisons as described in the figure legends. Mtb fold change (Fig. 1G) analyzed by unpaired t-test. All statistical tests were run using GraphPad Prism 8. *. P values ≤ 0.05 were considered significant. P values were conventionally denoted in the figures as *P ≤ 0.05; **P ≤ 0.01; ***P ≤ 0.001.* \*\*\*\**P* ≤ 0.0001. All error bars represent standard deviation and center bars indicate means.

### Study approval

All experiments were performed with the full knowledge and approval of the Standing Institutional Animal Care and Use Committee (IACUC) and Institutional Biosafety Committee (IBC) at Boston University (IACUC protocol PROTO201800218, IBC protocol 19-875) in accordance with relevant guidelines and regulations.

## Acknowledgement

This work was sponsored by R01 HL133190 and R01 HL126066 to IK and WB. BNK and OSR were supported by NIH/NCI grant R01CA244660. The authors are grateful to Drs. John Connor, Rahm Gummuluru, Andrew Henderson, Alla Grishok and Ben Wolozin for helpful discussions.

## Author contributions

EB, SMY, LK, WB and IK designed research studies;

EB, SMY, KS, BB, SC, HAC NS conducted experiments and performed data acquisition;

VZ, OR, AG, BNK, NC, EB, SMY, KS analyzed the data and prepared figures and publication materials;

EB, SY, VZ, AG, LK and IK wrote the manuscript.

## Notes

The authors have declared that no conflict of interest exists.

### Competing Interest Statement

The authors have declared no competing interest.

