## Supplemental Tables for "Maladaptive oxidative stress cascade drives type I interferon hyperactivity in TNF activated macrophages promoting necrosis in murine tuberculosis granulomas"

**Supplemental Table 1:** Significance values for comparisons in Figure 2A and 3A. Samples numerically labeled according to diagram in Figure 2A. Significance values by Two-way ANOVA. *$p < 0.05$, ** $p < 0.01$, *** $p < 0.001$, **** $p < 0.0001$.

| | IFN | Rsad2 | iNOS | IL1Ra | Trib3 | Chac1 |
|---|---|---|---|---|---|---|
| 1 vs 2 | **** | **** | **** | **** | **** | **** |
| 1 vs 3 | ns | *** | **** | ns | ns | ns |
| 1 vs 5 | ns | *** | **** | ** | ns | ns |
| 2 vs 4 | **** | **** | **** | ** | ** | **** |
| 2 vs 6 | **** | **** | **** | **** | **** | **** |
| 3 vs 4 | ns | **** | **** | **** | **** | * |
| 3 vs 5 | ns | ns | ns | *** | ns | ns |
| 4 vs 6 | ns | **** | **** | **** | **** | ns |
| 5 vs 6 | ns | ns | ns | **** | * | ns |

**Supplemental Table 2.** Gene set enrichment analysis results for the top enriched gene sets from the KEGG, MSigDB Hallmark, Reactome databases when comparing B6.Sst1S vs B6.WT BMDMs stimulated with TNF for 12 h. Output includes gene set names and FDR corrected p-values. The analysis was performed using FPKM data and the camera function from the limma R package.

### Gene Set Enrichment Analysis

| | Gene set | FDR |
|---|---|---|
| **Overrepresented genesets for comparison B6.Sst1S.TNF vs B6.TNF** | SUMOYLATION (Reactome) | 0.002714 |
| | SUMOYLATION OF DNA DAMAGE RESPONSE AND REPAIR PROTEINS (Reactome) | 0.002778 |
| | TNFA SIGNALING VIA NFKB (MSigDB Hallmark) | 0.002934 |
| | SPLICEOSOME (KEGG) | 0.00816 |
| | DNA DOUBLE-STRAND BREAK REPAIR (Reactome) | 0.016533 |
| | DNA REPAIR (Reactome) | 0.034949 |
| | HOMOLOGOUS RECOMBINATION (KEGG) | 0.037469 |
| | E2F TARGETS (MSigDB Hallmark) | 0.054885 |
| **Underrepresented genesets for comparison B6.Sst1S.TNF vs B6.TNF** | LYSOSOME (KEGG) | 8.26E-09 |
| | OXIDATIVE PHOSPHORYLATION (KEGG) | 1.36E-05 |
| | OXIDATIVE PHOSPHORYLATION (MSigDB Hallmark) | 6.82E-05 |
| | NONSENSE MEDIATED DECAY (NMD) INDEPENDENT OF THE EXON JUNCTION COMPLEX (EJC) (Reactome) | 0.00027 |
| | FATTY ACID METABOLISM (MSigDB Hallmark) | 0.002066 |
| | RESPONSE TO ELEVATED PLATELET CYTOSOLIC CA2+ (Reactome) | 0.002126 |
| | DETOXIFICATION OF REACTIVE OXYGEN SPECIES (Reactome) | 0.002257 |
| | FATTY ACID METABOLISM (KEGG) | 0.002903 |
| | FERROPTOSIS (KEGG) | 0.004529 |
| | MTORC1 SIGNALING (MSigDB Hallmark) | 0.00642 |
| | REACTIVE OXYGEN SPECIES PATHWAY (MSigDB Hallmark) | 0.008065 |
| | PEROXISOME (MSigDB Hallmark) | 0.014752 |
| | CHOLESTEROL_HOMEOSTASIS (MSigDB Hallmark) | 0.030761 |

**Supplemental Table 3.** The functional profiling and transcription factor binding sites analysis (TFBs) was performed for two sets of genes which represent B6.Sst1S and B6.WT BMDMs responses to TNF.

### Functional pathway profiling

| | Enriched pathways | Adjusted p.value | genes |
|---|---|---|---|
| Sst1.S response | TNF signaling pathway (KEGG) | 0.0002965 | Map3K5, Casp3, Jun, Nfkb1, Mapk8, Edn1, Ripk1, Il6 |
| | Cellular Senescence (Reactome) | 0.0040589 | Map3K5, Jun, Cbx8, Mapk8, Mapk7, Mdm2, Ets1, Kdm6B |
| | MyD88-independent TLR4 cascade (Reactome) | 0.0065647 | Jun, Nfkb1, Mapk8, Tlr4, Ripk1, Mapk7 |
| | Toll Like Receptor 4 (TLR4) Cascade (Reactome) | 0.0129301 | Jun, Nfkb1, Mapk8, Tlr4, Ripk1, Mapk7 |
| | Oxidative Stress Induced Senescence (Reactome) | 0.0247333 | Map3K5, Jun, Cbx8, Mapk8, Mdm2, Kdm6B |
| B6 response | Detoxification of Reactive Oxygen Species (Reactome) | 1.48E-15 | Gpx1, Txn2, Prdx1, Gpx3, Cat, Txnrd1, Prdx2, Sod2, Sod1, Prdx3, Prdx5, Gsr |
| | TP53 Regulates Metabolic Genes (Reactome) | 3.4733E-05 | Akt1, Prdx1, G6Pdx, Txnrd1, Prdx2, Prdx5, Gsr |
| | Ferroptosis (KEGG) | 9.9415E-05 | Slc11A2, Gpx4, Slc7A11, Atg7, Prnp, Gclm |
| | HIF-1 signaling pathway (KEGG) | 0.003628 | Akt1, Mapk3, Ldha, Egln1, Pdk1, Mapk1, Camk2G |
| | Peroxisome (KEGG) | 0.0054955 | Cat, Mpv17, Idh1, Sod2, Sod1, Prdx5 |

**Supplemental Table 4.** The table shows the identified list of transcription factors associated with differences between activated genes in response to TNF stimulation in B6.Sst1S vs B6.WT BMDMs.

### Transcription factor binding sites analysis

**B6.sst1S response**

| Motif ID | Possible TF | NES |
|---|---|---|
| transfac_M04823 | E2f4 | 4.46 |
| transfac_M04797 | Egr1 | 4.33 |
| transfac_M04683 | Nfya | 4.09 |
| taipale_TEAD4 | Tead4,Cebpd | 4.05 |
| transfac_M07390 | Nfe2l1 | 3.96 |
| transfac_M02743 | E2f3 | 3.94 |
| cisbp_M2272 | E2f4 | 3.91 |
| factorbook_NFY | Nfyb,Nfya | 3.90 |
| cisbp_M4473 | Pbx3 | 3.87 |
| cisbp_M4504 | Egr1 | 3.75 |

**B6.WT response**

| Motif ID | Possible TF | NES |
|---|---|---|
| cisbp_M6156 | Bach1 | 5.72 |
| cisbp_M1966 | Nfe2l2 | 5.48 |
| factorbook_NFE2 | Nfe2 | 5.46 |
| jaspar_MA0150.2 | Nfe2l2 | 5.43 |
| hocomoco_BACH1 | Bach1 | 5.23 |
| taipale_ELK1 | Elk1 | 4.52 |
| jaspar_MA0501.1 | Nfe2 | 4.48 |
| transfac_M02022 | Mafk | 4.32 |
| cisbp_4629 | Nfe2 | 4.28 |
| transfac_M07374 | Bach1 | 4.12 |

**Supplemental Table 5.** Master regulator analysis was performed using Virtual Inference of Protein Activity by Enriched Regulon Analysis (VIPER) algorithm, the mice macrophage gene regulatory network and log2 transformed FPKM values of genes from gene ontology category GO: 0006979. The table shows the identified list of transcription factors associated with differences between activated genes in response to TNF stimulation in B6.Sst1S vs B6.WT BMDMs.

### Master regulator analysis

| TFs | t | p.value |
|---|---|---|
| Nfe2l1 | 12.34 | 0.000248 |
| Atf5 | 9.49 | 0.000688 |
| E2f3 | 4.82 | 0.00855 |
| E2f6 | -10.51 | 0.000463 |
| Zfp523 | -10.76 | 0.000423 |
| Zfp623 | -12.46 | 0.000239 |
| Zkscan8 | -13.39 | 0.00018 |
| Zfp568 | -16.23 | 8.44E-05 |
| Zfp251 | -58.59 | 5.08E-07 |

**Supplemental Table 6.**

| Regulon | Size | NES | p.value | FDR |
|---|---|---|---|---|
| Zfp445 | 212 | 2.92 | 3.54E-03 | 0.0424 |
| Stat1 | 81 | 2.89 | 3.90E-03 | 0.0424 |
| Adnp2 | 938 | 2.72 | 6.57E-03 | 0.0579 |
| Irf7 | 51 | 2.71 | 6.69E-03 | 0.0579 |
| Zfp777 | 525 | 2.71 | 6.77E-03 | 0.0579 |
| Mbd3 | 36 | -2.86 | 4.26E-03 | 0.0429 |
| Rit1 | 42 | -2.88 | 3.97E-03 | 0.0424 |
| Thap7 | 65 | -2.88 | 3.92E-03 | 0.0424 |
| Foxk2 | 291 | -2.95 | 3.16E-03 | 0.0424 |
| Gtf3a | 51 | -2.97 | 3.01E-03 | 0.0424 |
| Thyn1 | 81 | -3.2 | 1.40E-03 | 0.0239 |
| Spi1 | 94 | -3.31 | 9.34E-04 | 0.0177 |
| Mlx | 74 | -3.35 | 7.99E-04 | 0.0171 |
| Tef | 468 | -3.76 | 1.70E-04 | 0.00415 |
| Max | 396 | -4.02 | 5.85E-05 | 0.00167 |
| Zfp664 | 88 | -4.22 | 2.49E-05 | 0.00085 |
| Ssrp1 | 114 | -4.25 | 2.10E-05 | 0.00085 |
| E2f6 | 832 | -4.31 | 1.64E-05 | 0.00085 |
| Mta2 | 131 | -4.81 | 1.54E-06 | 0.00013 |
| Nfyc | 99 | -4.88 | 1.09E-06 | 0.00013 |

**Supplemental Table 7:** Sst1S BMDMs treated with Deferoxamine as described in Figure 5C, and harvested for qRT-PCR analysis for IFNβ. Values represent fold change in respective RNA compared to untreated control. Standard deviations and p-values for individual experiment were calculated based on technical replicates.

| IFN | Expt | TNF | Standard Deviation TNF | TNF + DFOM | Standard deviation DFOM | p value |
|---|---|---|---|---|---|---|
| | 1 | 13.67 | 1.47 | 1.46 | 0.12 | 0.0046 |
| | 2 | 9.15 | 0.74 | 3.64 | 0.77 | 0.0009 |
| | 3 | 5.19 | 0.49 | 3.89 | 0.51 | 0.0335 |

**Supplemental Table 8:** Sst1S BMDMs treated with Deferoxamine as described in Figure 5C, and harvested for qRT-PCR analysis for Trib3. Values represent fold change in respective RNA compared to untreated control. Standard deviations and p-values for individual experiment were calculated based on technical replicates.

| Trib3 | Expt | TNF | Standard Deviation TNF | TNF + DFOM | Standard deviation DFOM | p value |
|---|---|---|---|---|---|---|
| | 1 | 8.43 | 0.24 | 3.95 | 0.68 | 0.0038 |
| | 2 | 24.45 | 0.84 | 18.23 | 3.3 | 0.0744 |
| | 3 | 17.5 | 1.63 | 13.47 | 1.14 | 0.0296 |

**Supplemental Table 9:** Sst1S BMDMs treated with Ferrostatin as described in Figure 5D, and harvested for qRT-PCR analysis for IFNβ. Values represent fold change in respective RNA compared to untreated control. Standard deviations and p-values for individual experiment were calculated based on technical replicates.

| IFN | Expt | TNF | Standard Deviation TNF | TNF + Fer1 | Standard deviation Fer1 | p value |
|---|---|---|---|---|---|---|
| | 1 | 10.99 | 2.11 | 7.28 | 0.25 | 0.039 |
| | 2 | 13.67 | 1.47 | 3.17 | 2.08 | 0.002 |
| | 3 | 9.15 | 0.74 | 5.24 | 0.94 | 0.005 |

**Supplemental Table 10:** Sst1S BMDMs treated with Ferrostatin as described in Figure 5D, and harvested for qRT-PCR analysis for Trib3. Values represent fold change in respective RNA compared to untreated control. Standard deviations and p-values for individual experiment were calculated based on technical replicates.

| Trib3 | Expt | TNF | Standard Deviation TNF | TNF + Fer1 | Standard deviation Fer1 | p value |
|---|---|---|---|---|---|---|
| | 1 | 8.43 | 0.24 | 3.69 | 0.4 | 0.0002 |
| | 2 | 24.45 | 0.84 | 18.6 | 0.58 | 0.001 |
| | 3 | 17.5 | 1.63 | 12.89 | 1.78 | 0.03 |

**Supplemental Table 11:** OCR and ECAR data for experiments described in Figure 6B. Oligomycin added before measurement 6, FCCP added before measurement 9, Antimycin added before measurement 12. Each measurement represents the mean of 6 technical replicates for the time of measurement.

| | | | Measurements | | | | | | | | | | | | | | | |
|---|---|---|---|---|---|---|---|---|---|---|---|---|---|---|---|---|---|---|
| | | Sample | 1 | 2 | 3 | 4 | 5 | 6 | 7 | 8 | 9 | 10 | 11 | 12 | 13 | 14 | 15 | 16 |
| Expt #1 | OCR | WT Ctrl | 105.23 | 95.50 | 92.53 | 92.18 | 92.02 | 40.62 | 39.83 | 38.54 | 147.99 | 123.34 | 114.20 | 28.35 | 26.15 | 26.19 | 25.62 | 24.47 |
| | | WT TNF | 149.37 | 138.54 | 135.37 | 136.35 | 137.55 | 53.93 | 50.06 | 48.91 | 145.28 | 137.77 | 137.44 | 27.67 | 28.54 | 29.13 | 28.86 | 28.06 |
| | | Sst1-S Ctrl | 104.37 | 94.85 | 92.14 | 92.37 | 92.72 | 36.58 | 36.74 | 35.94 | 161.23 | 131.51 | 118.87 | 19.56 | 21.05 | 22.77 | 23.05 | 22.71 |
| | | Sst1-S TNF | 114.94 | 107.01 | 104.30 | 104.79 | 105.15 | 43.44 | 42.93 | 42.46 | 115.59 | 102.66 | 98.28 | 25.86 | 26.88 | 26.80 | 26.60 | 25.86 |
| | ECAR | WT Ctrl | 31.82 | 19.99 | 21.19 | 22.73 | 23.77 | 47.75 | 28.46 | 30.04 | 54.11 | 48.16 | 46.99 | 41.39 | 39.53 | 38.63 | 38.30 | 38.66 |
| | | WT TNF | 48.35 | 33.79 | 35.17 | 37.35 | 38.69 | 63.83 | 72.34 | 80.04 | 101.89 | 104.45 | 101.23 | 98.86 | 97.14 | 93.87 | 92.58 | 92.01 |
| | | Sst1-S Ctrl | 33.34 | 23.17 | 23.71 | 25.02 | 26.33 | 44.78 | 26.81 | 27.75 | 54.25 | 47.60 | 47.34 | 41.37 | 39.53 | 38.00 | 38.84 | 39.27 |
| | | Sst1-S TNF | 44.63 | 34.16 | 36.35 | 39.42 | 41.61 | 62.67 | 66.45 | 71.03 | 83.58 | 85.10 | 86.19 | 87.55 | 83.61 | 81.40 | 81.13 | 81.37 |
| Expt #2 | OCR | WT Ctrl | 80.92 | 77.14 | 75.50 | 75.03 | 75.26 | 30.88 | 30.29 | 30.37 | 49.95 | 50.55 | 48.89 | 21.67 | 21.57 | 20.99 | 21.16 | 20.52 |
| | | WT TNF | 150.92 | 145.14 | 143.08 | 142.78 | 142.81 | 57.77 | 56.57 | 56.36 | 89.59 | 84.49 | 82.93 | 37.75 | 38.85 | 38.09 | 37.20 | 36.12 |
| | | Sst1-S Ctrl | 48.77 | 45.87 | 45.37 | 45.29 | 44.79 | 28.52 | 28.13 | 27.83 | 39.49 | 40.27 | 38.88 | 23.55 | 23.01 | 21.86 | 21.45 | 22.40 |
| | | Sst1-S TNF | 77.14 | 72.92 | 72.23 | 71.20 | 71.11 | 40.57 | 40.05 | 39.58 | 53.82 | 51.01 | 49.93 | 33.22 | 32.11 | 31.28 | 31.36 | 30.75 |
| | ECAR | WT Ctrl | 35.27 | 35.26 | 36.58 | 37.04 | 37.71 | 44.32 | 20.73 | 17.47 | 13.11 | 7.76 | 7.17 | 3.49 | 1.77 | 1.56 | 1.58 | 1.28 |
| | | WT TNF | 69.87 | 62.64 | 69.06 | 72.85 | 74.97 | 56.94 | 48.77 | 56.57 | 42.40 | 41.98 | 45.90 | 22.25 | 23.59 | 25.85 | 27.32 | 28.31 |
| | | Sst1-S Ctrl | 28.13 | 22.33 | 24.79 | 26.38 | 27.34 | 41.24 | 21.03 | 17.08 | 10.67 | 5.00 | 4.16 | 0.04 | -1.07 | -0.81 | -0.70 | -0.52 |
| | | Sst1-S TNF | 38.07 | 33.87 | 38.56 | 41.77 | 44.08 | 52.64 | 38.58 | 38.85 | 23.84 | 19.77 | 20.93 | 10.13 | 9.22 | 10.38 | 11.10 | 11.34 |
| Expt #3 | OCR | WT Ctrl | 74.68 | 68.81 | 65.75 | 64.03 | 63.23 | 31.80 | 31.36 | 31.52 | 74.93 | 51.34 | 41.92 | 21.64 | 21.55 | 21.46 | 20.77 | 20.72 |
| | | WT TNF | 130.72 | 119.80 | 114.46 | 112.46 | 111.28 | 51.72 | 50.07 | 49.33 | 65.06 | 58.39 | 54.33 | 29.53 | 29.21 | 28.71 | 27.63 | 27.34 |
| | | Sst1-S Ctrl | 54.53 | 50.32 | 48.42 | 47.81 | 47.06 | 24.16 | 24.31 | 23.75 | 46.63 | 39.90 | 32.84 | 17.68 | 16.86 | 16.73 | 16.30 | 15.32 |
| | | Sst1-S TNF | 53.56 | 51.57 | 48.31 | 48.04 | 47.72 | 30.36 | 27.65 | 27.86 | 32.29 | 30.08 | 29.76 | 20.54 | 19.51 | 19.35 | 19.09 | 18.41 |
| | ECAR | WT Ctrl | 39.97 | 13.86 | 10.73 | 11.52 | 11.94 | 13.13 | 7.30 | 8.01 | 5.32 | 7.05 | 10.29 | 11.53 | 9.49 | 8.88 | 9.02 | 8.41 |
| | | WT TNF | 61.75 | 25.77 | 21.46 | 22.19 | 22.76 | 19.58 | 16.88 | 17.83 | -9.95 | 12.17 | 18.33 | 17.76 | 17.07 | 16.39 | 16.56 | 16.23 |
| | | Sst1-S Ctrl | 34.15 | 8.99 | 5.28 | 5.93 | 6.62 | 5.97 | 5.25 | 5.82 | 1.47 | 1.63 | 6.38 | 8.34 | 6.34 | 5.41 | 6.14 | 5.98 |
| | | Sst1-S TNF | 51.51 | 21.42 | 17.50 | 18.11 | 18.56 | 16.88 | 16.66 | 17.79 | 5.99 | 16.38 | 20.12 | 16.98 | 17.20 | 17.60 | 18.41 | 18.26 |

**Supplemental Table 12:** Quantification of morphological changes observed by TEM, as described in Figure 6C. Images scanned were all at 50,000x resolution, as the representative image. Chi-square analysis reveals no difference between WT and Sst1S controls, but demonstrates significance between WT and Sst1S samples treated with TNF ($p = 0.0015$).

| | Images scanned | Images with Widened cristae | Images with voids | Total Mitochondria counted | Total w/ widened cristae | Total with voids |
|---|---|---|---|---|---|---|
| WT Ctrl | 8 | 3 | 0 | 64 | 3 (4.7%) | 0 (0%) |
| Sst1S Ctrl | 11 | 4 | 0 | 66 | 6 (9.1%) | 0 (0%) |
| WT TNF | 10 | 9 | 3 | 67 | 31 (46.2%) | 4 (6.0%) |
| Sst1S TNF | 12 | 12 | 5 | 98 | 65 (66.3%) | 12 (12.2%) |

**Supplemental Table 13**

RNA primer sequences.

| Gene | Primer forward | Primer reverse |
|---|---|---|
| 18S | TCAAGAACGAAAGTCGGAGGT | CGGGTCATGGGAATAACG |
| IFNb | ATGAGTGGTGGTTGCAGGC | TGACCTTTCAAATGCAGTAGATTC |
| Trib3 | GCAAAGCGGCTGATGTCTG | AGAGTCGTGGAATGGGTATCTG |
| Chac1 | CCTGCTACCCTGCTC TTACCT | GAGCTTGGCTCCTCAGGTC |
| IL1ra | CGCCCTTCTGGGAAAAGACC | CCGTGGATGCCCAAGAACAC |
| iNOS | Kramnik-Lab | Kramnik-Lab |
| Hspa1a | GATTTGTTTTGCAGGACAGC | GGGGAGAGTCCAAACACAAA |
| Rsad2 | AAGCTGAGGAGGTGGTGCAG | GAAAACCTTCCAGCGCACAG |
