## Supplemental Figures for "Maladaptive oxidative stress cascade drives type I interferon hyperactivity in TNF activated macrophages promoting necrosis in murine tuberculosis granulomas"

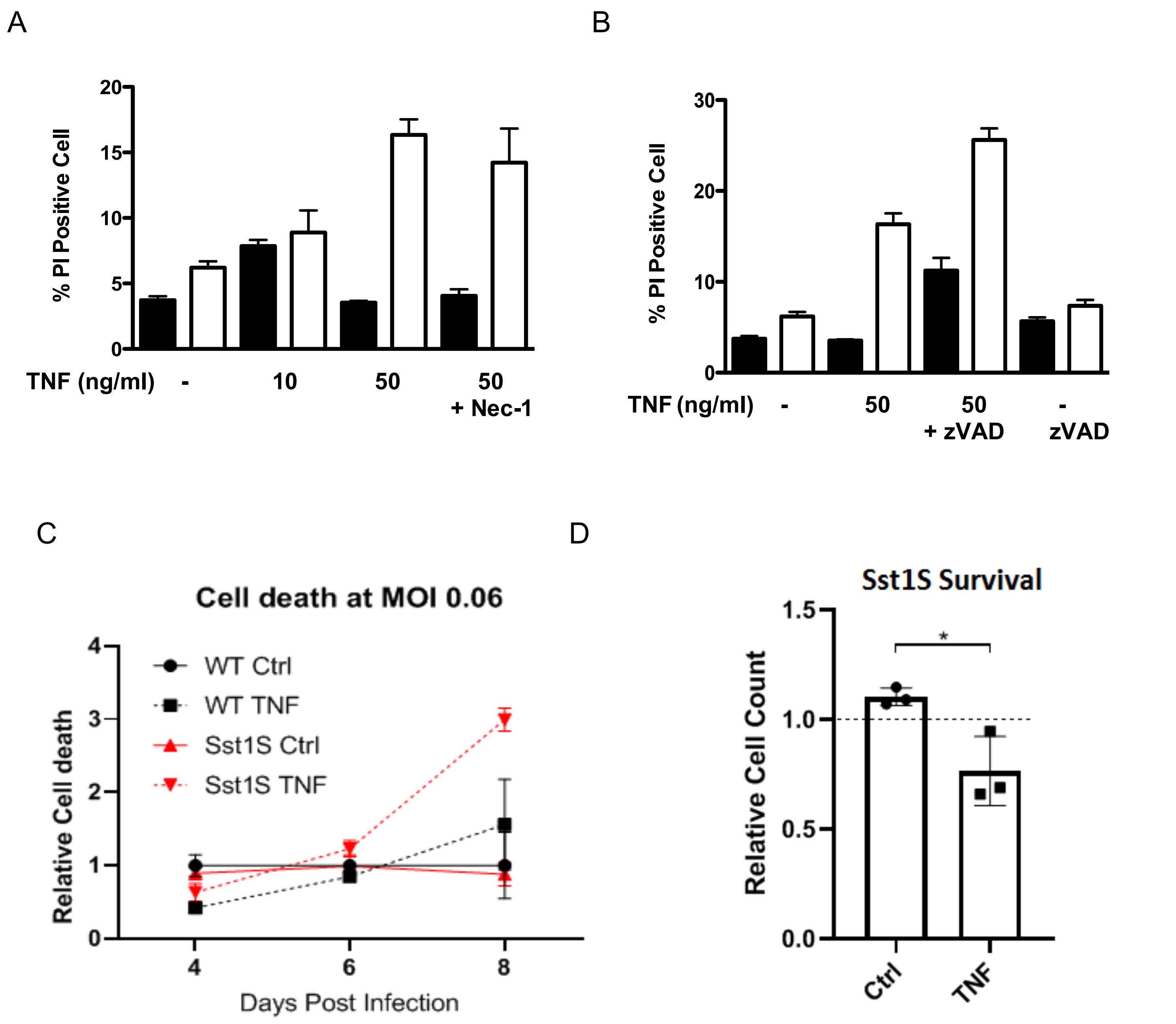


**Supplemental Figure 1.**

**A - B.** Increased cell death of B6.Sst1S BMDMs after 48 h of TNF stimulation was not inhibited by Nec1 (A) or zVAD (B).

**C – D**. Cell death of BMDMs during chronic infection with virulent *M.tb* in vitro is mediated by TNF in sst1-dependent manner. The Sst1S and WT BMDMs were infected with *M.tb* H37Rv (MOI 0.1) *in vitro*. 10 ng/mL TNF was added to experimental group samples 24h before infection, and was added again immediately following media changes, which occurred every 2 days. At harvest, infected cells were stained with “Live or Dye” fixable viability stain (Biotium), fixed in 4% PFA, co-stained with Hoechst, and counted using the Nexelcom Celigo fluorescent plate reader microscope.
**C.** Representative time-course; relative cell death calculated using total dead cells divided by total cells counted, normalized to WT untreated. Error bars indicate standard deviation of technical replicates.
**D.** Repeat experiments at Day 8 post-infection. Relative cell count determined as total cells minus dead cells, normalized to Sst1S uninfected controls. Error indicate standard deviation of three experiments. Significance by paired T-test.

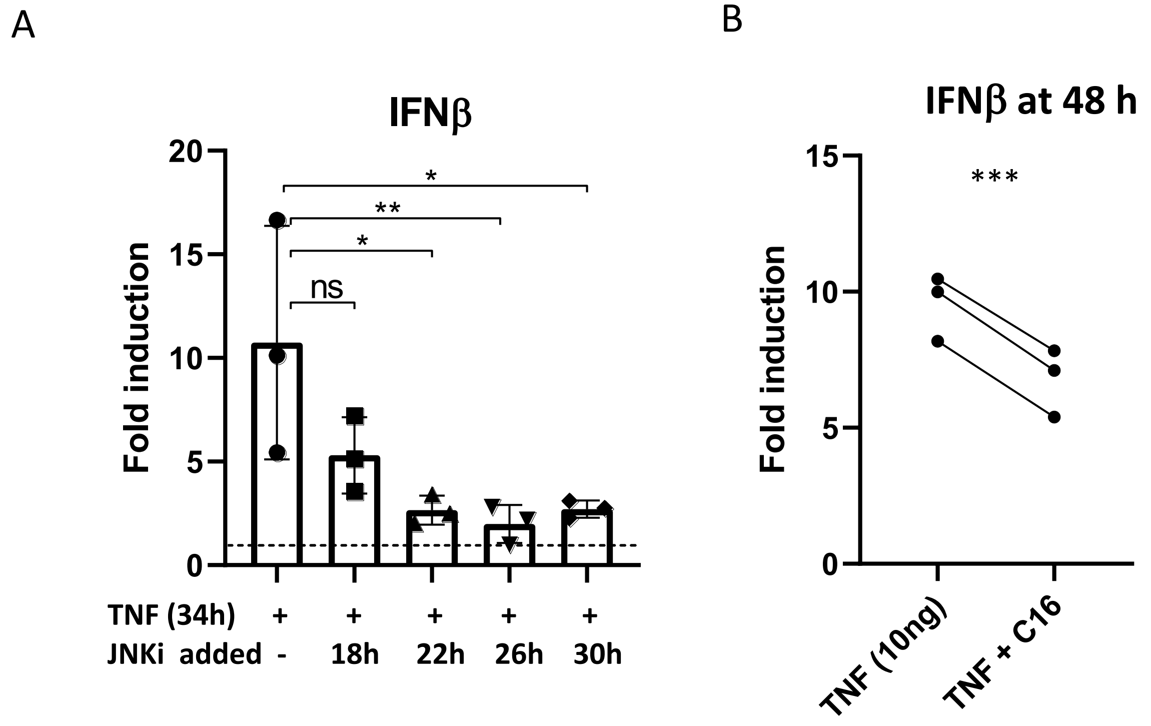


**Supplemental Figure 2.**

**A.** JNK inhibitor reduces levels of IFNβ transcript induced by TNF. Sst1S BMDMs were stimulated with TNF for 34 hours, or left unstimulated. JNK inhibitor was added to samples 18, 22, 26, or 30 hours after TNF, and RNA samples were collected at 34 h. The IFNβ mRNA was measured using qRT-PCR. Fold induction is normalized to untreated controls. Single experiment shown; error bars represent standard deviation of technical replicates, and significance values and error bars are based on technical replicates.

**B.** Sst1S BMDMs were treated with TNF, and analyzed for IFNb mRNA expression, as in Fig.3F. Significance by paired T-test, *** $p<0.001$

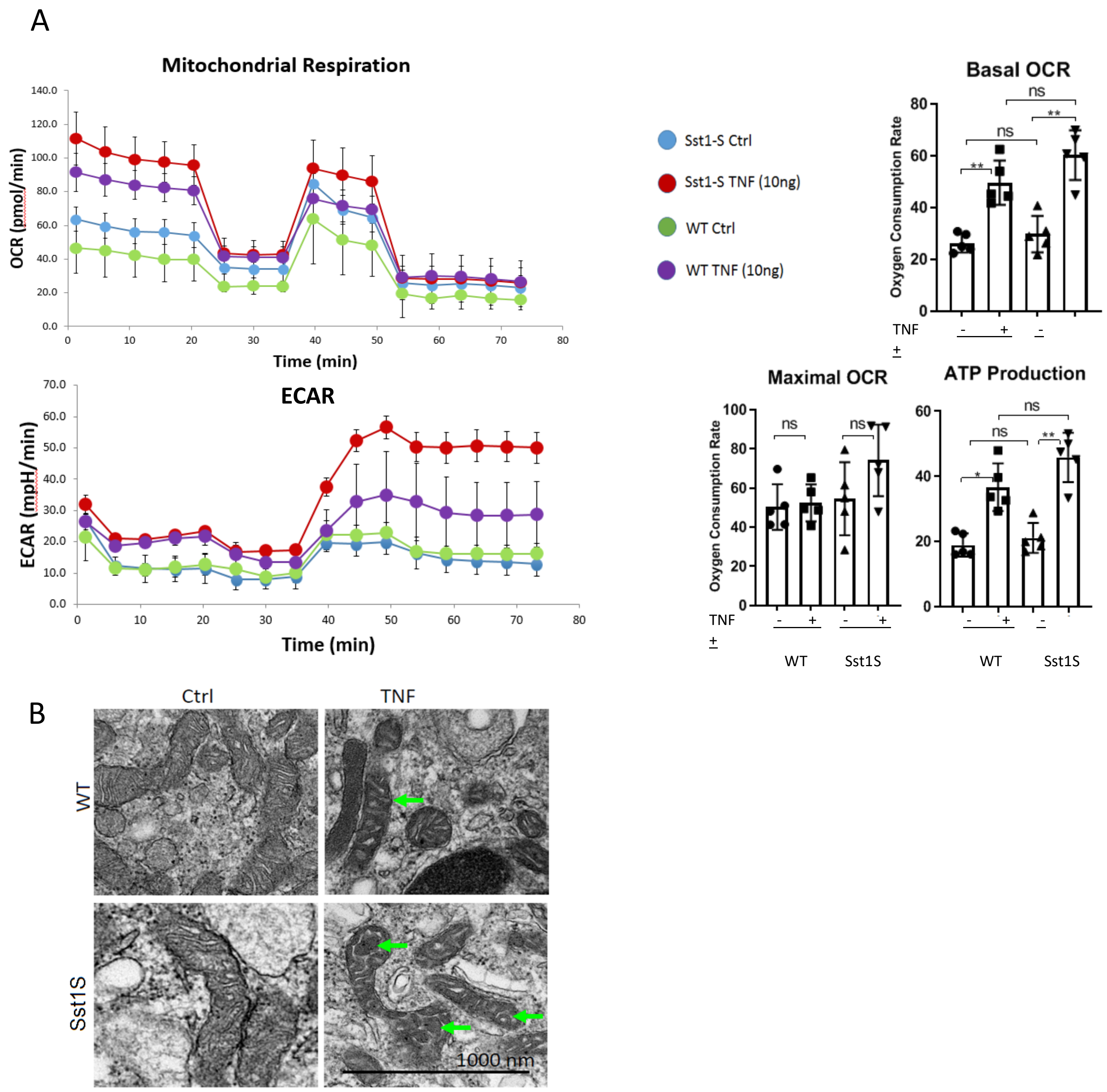


**Supplemental Figure 3.**
**A**. WT and Sst1S BMDMs were isolated and treated with 10ng/mL TNF or left untreated. 18 hours after stimulation, cell media was changed to Agilent Seahorse medium, supplemented with glucose, pyruvate and L-glutamine, and then assayed using the Agilent Seahorse Extracellular Flux Analyzer, using standard protocols for Oxygen Consumption Rate analysis.
Left panels: Representative experiment: traces for OCR (upper panel) and ECAR (lower panel). Right panels: Graphical analysis of Basal OCR, Maximal OCR, and ATP production for each strain. N=5 experiments, error bars represent standard deviation. Statistical analysis by RM-One way ANOVA with multiple comparisons. *$p < 0.05$, ** $p < 0.01$.

**B**. WT and Sst1S BMDMs were treated with 10ng/mL TNF for 48h, or left untreated (Ctrl). Cells were then collected and fixed for TEM imaging. 50000x TEM images (1000nm scale bar provided) reveals mitochondrial ultrastructure. Notable morphological changes observed include widened mitochondrial cristae in TNF-treated BMDMs, regardless of strain, and large mitochondrial matrix “voids” in Sst1S mitochondria. Both morphological phenomena indicated by green arrows. Additional quantification of images provided in supplemental table 12.

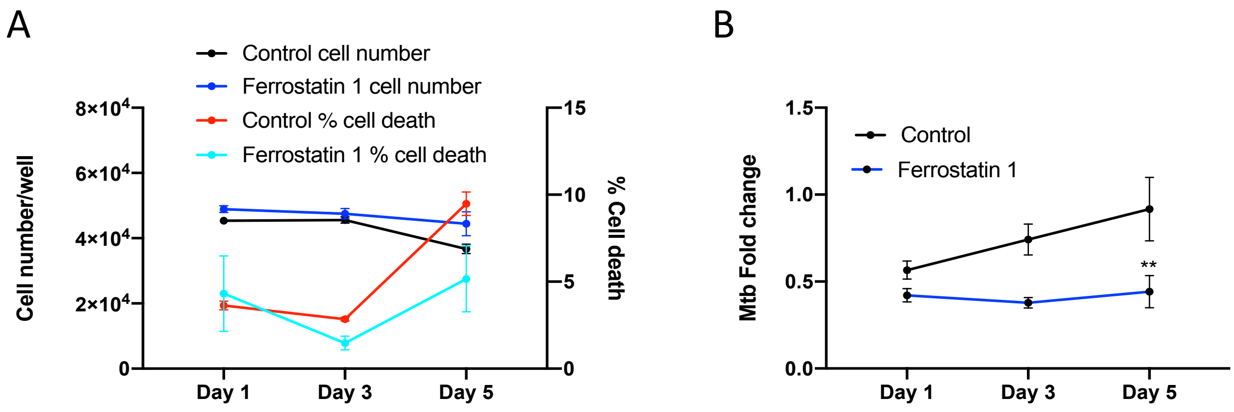


**Supplemental Figure 4.**

B6.Sst1S cells were pretreated with TNFa (10 ng/mL) for 8 h and subsequently treated with TNF in combination with Ferrostatin 1 (1 µM) for 16h.

**A**. The cells were survival and death at 1, 3, and 5 days post-infection.

**B**. The *M.tb* loads were quantified using a qPCR-based method, normalized to a BCG spike for internal control. Error bars indicate standard deviation of technical replicates, significance
